# Non-random segregation of mitochondria during asymmetric cell division contributes to cell fate divergence in daughter cells

**DOI:** 10.1101/2024.09.05.611553

**Authors:** Ioannis Segos, Jens Van Eeckhoven, Simon Berger, Nikhil Mishra, Eric J. Lambie, Barbara Conradt

## Abstract

The non-random segregation of organelles has been proposed to be an intrinsic mechanism that contributes to cell fate divergence during asymmetric cell division; however, *in vivo* evidence is sparse. Using super-resolution microscopy, we analysed the segregation of organelles during the division of the neuroblast QL.p in *C. elegans* larvae. QL.p divides to generate a daughter that survives, QL.pa, and a daughter that dies, QL.pp. We found that mitochondria segregate unequally by density and morphology and that this is dependent on mitochondrial fission and fusion. Furthermore, we found that mitochondrial density in QL.pp correlates with the time it takes QL.pp to die. We propose that low mitochondrial density in QL.pp promotes the cell death fate and ensures that QL.pp dies in a highly reproducible and timely manner. Our results provide the first *in vivo* evidence that the non-random segregation of mitochondria can contribute to cell fate divergence during asymmetric cell division.

## Introduction

The generation of cellular diversity via asymmetric cell division is of fundamental importance for (nearly) all (living) organisms^1,2^. It has been proposed that differential cell fate acquisition among the daughter cells is influenced by the non-random segregation of cellular organelles, particularly mitochondria^3–8^. Strong support for this idea has been obtained in studies of budding yeast, where mitochondria are segregated unequally between the mother cell and the bud. This leads to different mitochondrial functionalities in the two cells, resulting in the rejuvenation of the bud and replicative aging in the mother cell^9^. In this system, mitochondrial segregation has been shown to be actively controlled through myosin motor-dependent transport and Num1/NuMA-dependent anchoring^10–13^ to be disrupted by the absence of mitochondrial dynamics^9^. In the case of animal systems, there have been hints that asymmetric mitochondrial segregation might influence cell fate, e.g., we found that cells programmed to die during *C. elegans* development are characterized by fragmented mitochondria^14^. In mammalian cells, efforts have been made to establish links between mitochondria and asymmetric cell fate acquisition, but no common model has emerged^3,7,9,15–19^. Importantly, none of the studies in mammalian cells analysed mitochondria quantitatively and in real time as cells divide. Consequently, it has not been clearly established what the differences between mitochondria in daughter cells may be and how such differences might arise. In addition, whether differences between mitochondria contribute to differential cell fate acquisition among daughter cells remains an open question.

Here, we report how mitochondria segregate during an asymmetric cell division in *C. elegans* larvae. Specifically, we studied the asymmetric division of the neuroblast QL.p, which generates a smaller daughter (QL.pp) that adopts the ‘cell death’ fate and undergoes apoptosis, and a larger daughter (QL.pa) that survives and divides to form two cells (QL.paa and QL.pap), which differentiate into neurons (PVM and SDQL, respectively). We were able to follow mitochondrial segregation during QL.p division in real time and at super resolution. In addition to wild-type animals, we examined mutants in which either the asymmetry of QL.p division or mitochondrial dynamics are disrupted. Finally, by combining super-resolution imaging and long-term cell fate tracking, we discovered a positive correlation between mitochondrial density and the time it takes QL.pp to die. Our study represents the first quantitative and real time analysis of mitochondrial segregation in an asymmetrically dividing cell in a developing animal.

## Results

We utilized two complementary approaches to follow mitochondrial segregation during QL.p (and QL.pa) division in real time and at super resolution. In one set of experiments, we used a microfluidics-based immobilization system^20^ to image individual animals throughout the entire L1 stage of larval development (11-22 hours) and to acquire quantitative data on (i) mitochondrial segregation during QL.p division and (ii) mitochondrial segregation during QL.pa division (see **Fig.1a**). QL.p and QL.pa were tracked using an ultrafast confocal mode, and super-resolution imaging was limited to two time points during their division (metaphase and post-cytokinesis). 3D rendering was performed, and cellular and mitochondrial volumes determined using methodology that we recently established (Methods; DOI https://doi.org/10.21203/rs.3.rs-4320882/v1). In another set of experiments, we used nanobeads-based immobilization (DOI https://doi.org/10.21203/rs.3.rs-4320882/v1) to image individual animals at the L1 stage for 1-2 hours and to acquire quantitative data on mitochondrial segregation during QL.p division. To determine volume, morphology and position of individual mitochondria during QL.p division, we used super-resolution imaging at high temporal resolution (1 min intervals between metaphase and post-cytokinesis). Unless noted otherwise, 3D rendering was performed and cellular and mitochondrial volumes determined using methodology that we recently established (Methods; DOI https://doi.org/10.21203/rs.3.rs-4320882/v1).

**Figure 1.**
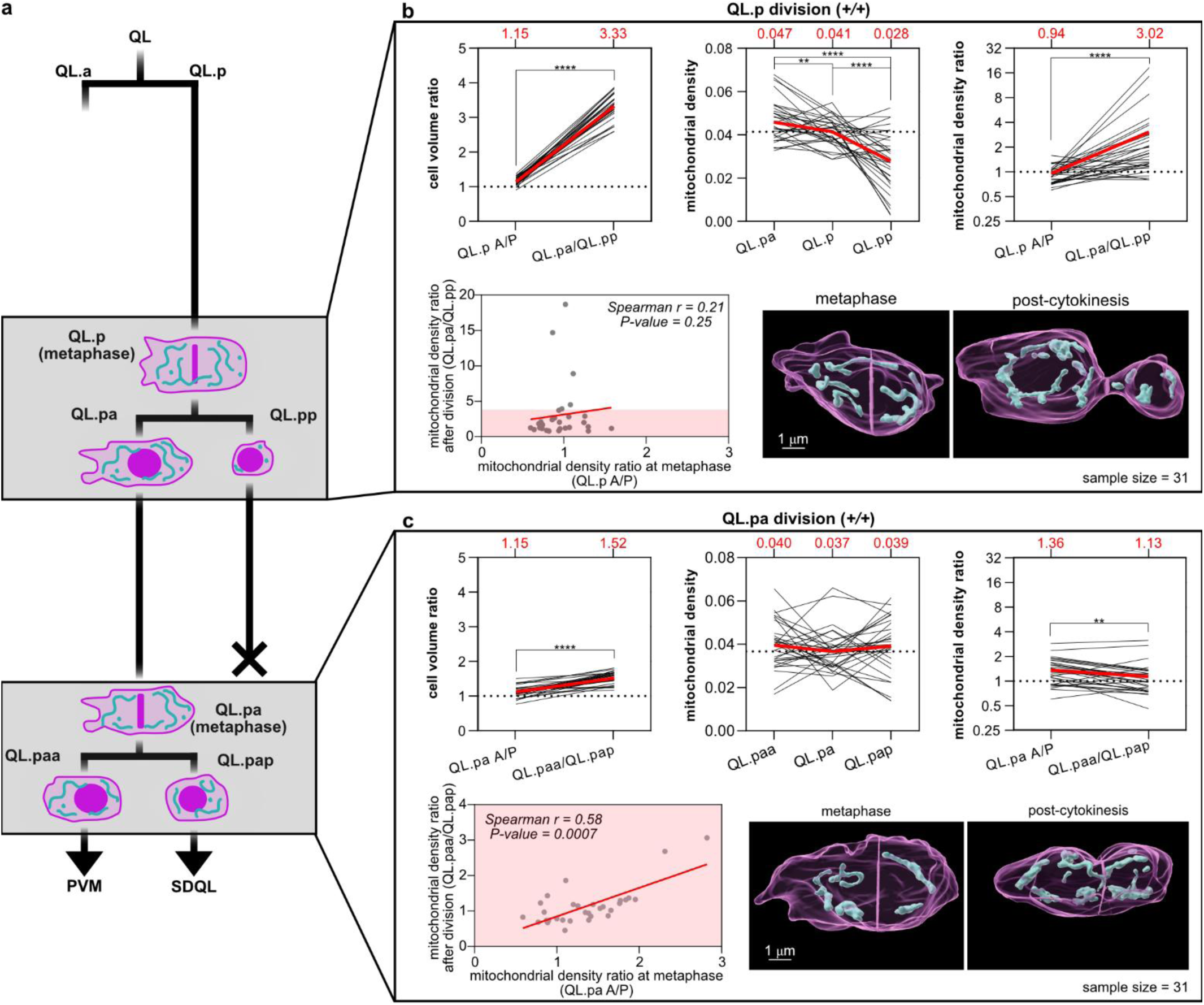
Mitochondrial partitioning is non-random and unpredictable during QL.p, but not QL.pa divisions. **a**) Schematics of the QL.p lineage recorded in wild-type animals expressing *bcIs153* transgene. The grey boxes highlight both QL.p and QL.pa divisions. **b**) Measurements of cell volume ratio, mitochondrial density and mitochondrial density ratio during QL.p division. Top left: cell volume ratio before (QL.p A/P sides) and after division (QL.pa/QL.pp); top centre: mitochondrial density before (QL.p) and after division (QL.pa and QL.pp); top right: mitochondrial density ratio before (QL.p A/P sides) and after division (QL.pa/QL.pp); bottom left: correlation between mitochondrial density ratio before (x axes) and after (y axes) division. **c**) Measurements of cell volume ratio, mitochondrial density and mitochondrial density ratio during QL.pa division. Top left: cell volume ratio before (QL.pa A/P sides) and after division (QL.paa/QL.pap); top centre: mitochondrial density before (QL.pa) and after division (QL.paa and QL.pap); top right: mitochondrial density ratio before (QL.pa A/P sides) and after division (QL.paa/QL.pap); bottom left: correlation between mitochondrial density ratio before (x axes) and after (y axes) division; bottom right: representative 3D volumes of QL.pa division. P values are calculated using the Wilcoxon matched pairs signed rank test (both b and c, top right), the paired t-test (both b and c, top left), the RM one-way ANOVA with the Benjamini, Krieger and Yekutieli correction (b, top centre), the Friedman test with the Benjamini, Krieger and Yekutieli correction (c, top centre), and the Spearman correlation (both b and c, bottom left). Normality was tested with the Shapiro-Wilk test. *: P value ≤ 0.05; **: P value ≤ 0.01; ***: P value ≤ 0.001; ****: P value ≤ 0.0001. In the top row plots of panels **a** and **b**, individual black lines represent the trends of each division between metaphase and post-cytokinesis or between QL.p and QL.pa and QL.pp. Red lines and red numbers = average (both b and c, top row). n=31 in a and b.

### Segregation of mitochondria is unequal by density during division of QL.p, but not QL.pa

We sought to determine how mitochondria segregate during QL.p division, which generates two daughter cells that adopt different fates, and during QL.pa division, which generates two neurons (**Fig.1a**, grey boxes). To quantitatively study these divisions, we used a strain in which cell membrane (myristoylated mCherry, magenta), chromatin (mCherry::histoneH1, magenta) and mitochondria (matrix-targeted GFP, cyan) are labelled specifically in Q lineage cells (*bcIs153* transgene). Imaging was performed using the microfluidics-based immobilization system^20^(see Methods).

At metaphase, the anterior (A) and posterior (P) halves of QL.p are essentially equal in volume (QL.p A/P volume ratio 1.15; **Fig.1b**, top left and bottom right). Post-cytokinesis, the two daughter cells are drastically unequal in volume, with a QL.pa/QL.pp volume ratio of 3.33 (**Fig.1b**, top left and bottom right). We found that mitochondria segregate unequally in terms of density (mitochondrial volume divided by cell volume) during QL.p division. Compared to the average mitochondrial density in QL.p, there is a significant increase in average mitochondrial density in the larger daughter QL.pa (from 0.041 to 0.047) and a corresponding decrease in the smaller daughter QL.pp (from 0.041 to 0.028) (**Fig.1b**, top centre). To correct for interindividual variance in the cellular and mitochondrial volume of the mother cell QL.p, we also determined the A/P ratios of average mitochondrial densities at metaphase (QL.p A/P) and post-cytokinesis (QL.pa/QL.pp) (**Fig.1b**, top right). We found that at metaphase, the QL.p A/P mitochondrial density ratio is 0.94, indicating that mitochondria are equally distributed prior to cell division. However, post-cytokinesis, the QL.pa/QL.pp mitochondrial density ratio was 3.02, indicating unequal mitochondrial distribution among the two daughter cells irrespective of their cell size difference. This suggests that mitochondria segregate unequally during QL.p division. To determine whether anterior-posterior mitochondrial distribution in QL.p is predictive of mitochondrial distribution in QL.pa and QL.pp, respectively, we plotted mitochondrial density ratios at metaphase against those at post-cytokinesis; however, we found no correlation (**Fig.1b**, bottom left; also see **FigS3**, *+/+*). Therefore, mitochondrial segregation during QL.p division is not preconfigured by the distribution of mitochondria at metaphase.

In the case of QL.pa, we found that at metaphase, the anterior and posterior halves of QL.pa are essentially equal in volume with a QL.pa A/P volume ratio of 1.15; post-cytokinesis, QL.paa is slightly larger than QL.pap with a QL.paa/QL.pap volume ratio of 1.52 (**Fig.1c**, top left and bottom right). We found that mitochondria are segregated equally during QL.pa division, as indicated by similar average mitochondrial densities in QL.pa and its daughter cells (0.037 vs 0.040 and 0.039), as well as similar average mitochondrial density ratios at metaphase and post-cytokinesis, which – in contrast to QL.p division - is significantly closer to 1 post-cytokinesis (1.13) (**Fig.1c**, top centre, top right, and bottom right). In contrast to QL.p division, we found a significant correlation between mitochondrial density ratios at metaphase (i.e. anterior-posterior mitochondrial distribution in QL.pa) and post-cytokinesis (i.e. mitochondrial distribution among the two daughter cells) (**Fig.1c**, bottom left). Therefore, during QL.pa division, mitochondria segregate equally, as preconfigured by their distribution at metaphase.

In summary, the segregation of mitochondria is unequal by density during QL.p division but not during QL.pa division. Moreover, the altered distribution of mitochondria after QL.p division suggests that mitochondrial segregation is a non-random actively controlled process.

### Unequal segregation during QL.p division may be restricted to mitochondria

Next, we wondered whether unequal segregation of organelles is a general feature of QL.p division. To answer this question, we generated strains in which cell membrane (myristoylated SFmTurquoise2ox, magenta), chromatin (SFmTurquoise2ox::his-24, magenta), mitochondria (*tomm-20*::mKate2 [in combination with lysosome labelling] or matrix-targeted GFP [in combination with ER labelling], cyan) and either lysosomes (eYFP::*cup-5*, green) (**Fig.S1a and S2**) or endoplasmic reticulum (ER) (SP12-mCherry-KDEL, green) (**Fig.S1b**) are labelled (*bcIs159* [lysosomes] and *bcIs160* [ER] transgenes). Imaging was performed at metaphase and post-cytokinesis using nanobead-based immobilization, and cellular size and organelle volumes were determined after 2D rendering (see Methods).

At metaphase, the distribution of lysosomes was slightly skewed towards the anterior side of QL.p, with an average QL.p A/P lysosome density ratio of 1.33 (**Fig. S1c**, left, QL.p A/P) but we observed a relatively symmetric distribution of ER (**Fig.S1d**, left). Post-cytokinesis, we observed essentially equal distribution of both lysosomes and ER with a subtle but significant increase in average density ratio between metaphase and post-cytokinesis in the case of ER (increase from 0.92 to 1.10) (**Fig. S1c,d**, left, QL.pa/QL.pp). Of note is that the average QL.pa/QL.pp mitochondrial density ratios observed using the nanobeads-based immobilization (2.02 and 1.89; **Fig.S1c,d**) are lower than those observed using the microfluidics-based immobilization system (3.02; **Fig.1b**). However, compared to their respective mitochondrial density ratios observed at metaphase (1.15 and 1.10; **Fig.S1c,d**), the QL.pa/QL.pp mitochondrial density ratios are still significantly increased after division. In this set of experiments, the anterior and posterior sides of QL.p at metaphase were essentially equal in size with QL.p A/P size ratios of 1.12 (lysosome) and 1.15 (ER). Post-cytokinesis, the two daughter cells generated were unequal in size with QL.pa/QL.pp size ratios of 3.33 (lysosomes) and 3.25 (ER) (**Fig. S1c, d,** right). These results suggest that lysosomes and ER are not unequally segregated during QL.p division. Therefore, unequal segregation during QL.p division may be restricted to mitochondria.

### Mitochondrial segregation during QL.p division is also unequal by morphology

To investigate the morphology and position of individual mitochondria during QL.p division, we used the same transgene as in **Fig.1** (*bcIs153*) in combination with nanobead-based immobilization and imaging at high temporal resolution (see Methods).

Using this experimental set-up, we observed essentially the same unequal segregation of mitochondria by density during QL.p division as described above using the microfluidics-based immobilization system (compare **Fig.1b** and **Fig.2a-d**, *+/+*). We quantified mitochondrial morphology by extracting mitochondrion-specific shape parameters and conducting a Principal Component Analysis (PCA). While mitochondria were morphologically comparable between the anterior and posterior sides of QL.p at metaphase, they differed between QL.pa and QL.pp post-cytokinesis. On average, mitochondria in QL.pa were larger and more elongated than in QL.pp, which inherited mostly fragmented mitochondria. This is evidenced by the significant difference along both PC1 (variance in surface area and volume of individual mitochondria or ‘size’) and PC3 (variance in sphericity and surface/volume ratio of individual mitochondria or ‘shape’) (**Fig.2b, e,** +/+). Thus, mitochondrial segregation is unequal not only in terms of density but also morphology.

**Figure 2.**
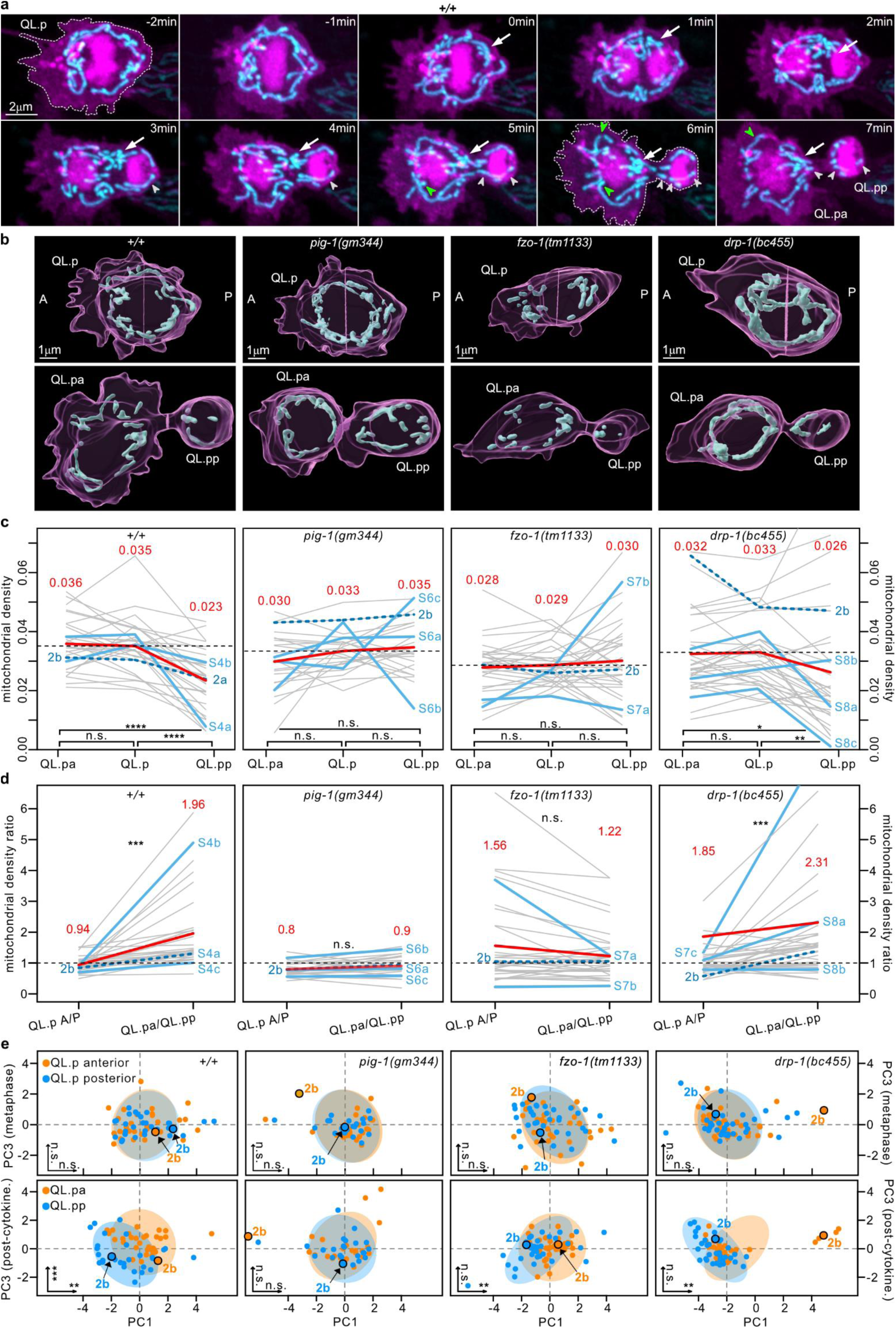
(previous page). Mitochondrial partitioning in *pig-1*, *fzo-1* and *drp-1* mutants is defective during QL.p division. **a**) Super-resolution live two-colour time series of QL.p division. Plasma membrane (myristoylated mCherry) and chromatin (mCherry::his-24) are showed in magenta, mitochondria (mtGFP) in cyan. Images are maximum intensity projections of aligned z-stacks. In all images, anterior is left and posterior is right. Arrows point to anteriorly directed transport of mitochondria. Green and white arrowheads point to mitochondrial fusion and fission, respectively. **b**) 3D rendering of representative samples of QL.p division at metaphase (top) and post-cytokinesis (bottom) in wildtype (*+/+*), *pig-1(gm344)*, *fzo-1(tm1133)*, and *drp-1(bc455)* (magenta: cell membrane; cyan: mitochondria). These examples refer to the blue lines in c and d and to the highlighted dots in e. **c**) mitochondrial density at metaphase (QL.p) and post-cytokinesis (QL.pa and QL.pp). Blue lines refer to the 3D rendered examples in b, whereas the green lines refer to timeseries illustrated in figures **S4**,**6,7,8**. Red lines and numbers = average. **d**) Mitochondrial density ratio at metaphase (QL.p A-side/P-side) and post-cytokinesis (QL.pa/QL.pp). Blue lines refer to the 3D rendered examples in b, whereas the green lines refer to timeseries illustrated in figures **S4**,**6,7,8**. Red lines and numbers = average. **e**) PCA analysis of mitochondrial morphological parameters at metaphase (top row) and post-cytokinesis (bottom row), with confidence ellipses for PC1 and PC3. Significance levels are indicated for both axes in the bottom left corner of the respective plots, with details on the analysis presented in the main text. In c P values are calculated using the RM one-way ANOVA or the Friedman test with the Benjamini, Krieger and Yekutieli correction. In d P values are calculated using the paired t-test or the Wilcoxon matched signed-rank test. Normality was tested using the Shapiro-Wilk test. *: P value ≤ 0.05; **: P value ≤ 0.01; ***: P value ≤ 0.001; ****: P value ≤ 0.0001. All data are from animals expressing *bcIs153* transgene.

The differences in mitochondrial morphology in QL.pa and QL.pp are also visually apparent in super-resolution time series (**Fig.2a and S4;** compare mitochondrial morphology in QL.p at metaphase/0min with mitochondrial morphologies in QL.pa and QL.pp post-cytokinesis/7min). These time series also revealed that between metaphase and post-cytokinesis, mitochondria are transported from the posterior to the anterior side (**Fig. 2a and S4**, white arrows). In addition, we observed mitochondrial fission and fusion events (**Fig. 2a and S4**, white and green arrowheads, respectively). Quantifying these events revealed that fission occurs significantly more frequently in the posterior side (**Fig. S5a, left**). Conversely, fusion occurs more often in the anterior side, but not significantly so (**Fig. S5a, right**). Therefore, during QL.p division, mitochondria are transported in a targeted way and mitochondrial fission occurs in a localized manner.

### Loss of QL.p’s ability to divide asymmetrically disrupts unequal mitochondrial segregation

Next, we asked whether the loss of QL.p’s ability to divide asymmetrically impacts unequal mitochondrial segregation. To address this, we used nanobead-based immobilization and imaging at high temporal resolution, and animals homozygous for a loss-of-function (lf) mutation of the gene *pig-1* (*pig-1*(*gm344*)).

*pig-1* encodes a PAR-1-like kinase that is orthologous to mammalian MELK (maternal embryonic leucine zipper kinase). When *pig-1* is inactivated, QL.p divides symmetrically and gives rise to two daughter cells of equal cell volume^21^ (**Fig. 2b**, *pig-1*(*gm344*)). We found that unequal mitochondrial segregation is severely affected in *pig-1*(lf) mutants: average mitochondrial densities in QL.p and its daughter cells are very similar (0.033 vs 0.030 and 0.035) and the average mitochondrial density ratios before and after division are essentially the same (0.8 and 0.9) (**Fig. 2c,d**). Morphological differences (PC1-size, PC3-shape) between QL.p daughter cells are lost in *pig-1*(lf) animals (**Fig. 2e**), and mitochondrial density ratios at metaphase correlate with mitochondrial density ratios post-cytokinesis (**Fig.S3**). This suggests that - unlike in wild type - mitochondria segregate equally, as preconfigured by their distribution at metaphase. Finally, in super-resolution time series, we observed that in *pig-1*(lf) animals, anteriorly directed mitochondrial transport is lost and mitochondrial fission and fusion events (white and green arrowheads) occur with similar frequencies in the posterior and anterior side of the cell (**Fig. S5b** and **S6a-c**). Overall, in the *pig-1*(lf) background, mitochondrial segregation during QL.p division becomes equal in terms of both density and morphology.

### Loss of mitochondrial dynamics compromises unequal mitochondrial segregation during QL.p division

Our finding that mitochondria divide and fuse during QL.p division prompted us to investigate the role of mitochondrial fission and fusion in unequal mitochondrial segregation. To that end, we used nanobead-based immobilization, imaging at high temporal resolution and animals homozygous for lf mutations of either *fzo-1* (*fzo-1(tm1133)*) or *drp-1* (*drp-1(bc455)*). *fzo-1* encodes the *C. elegans* ortholog of human Mitofusins 1 and 2, which are required for mitochondrial fusion, and its loss causes mitochondrial hyper-fission^22^. *drp-1* encodes the *C. elegans* ortholog of human Drp1, which is required for mitochondrial fission, and its loss causes mitochondrial hyper-fusion^23^. (The *drp-1* allele *bc455* was generated using CRISPR-Cas-based genome editing and is molecularly identical to the *drp-1* allele *tm1108* [see Methods].) In both mutants, QL.pa/QL.pp cell volume asymmetry is maintained (see **Fig.2B, Fig.S7 and Fig.S8**).

We found that in *fzo-1*(lf) mutants, average mitochondrial densities in QL.p and its daughter cells are similar (0.029 vs 0.028 and 0.030) (**Fig. 2c**, *fzo-1*(*tm1133*)), and average mitochondrial density ratios do not change significantly between metaphase and post-cytokinesis (1.56 and 1.22) (**Fig. 2d**). Mitochondria appear to be distributed haphazardly at metaphase, with more mitochondria located anterior or posterior of the metaphase plate, seemingly at random (**Fig. S7a-c**). The mitochondrial density ratios at metaphase are also highly predictive of the density ratios post-cytokinesis (**Fig. S3**). Moreover, compared to wild type, we observed fewer movements of mitochondria during anaphase and cytokinesis (**Fig. S7b**, see arrowheads pointing to posterior mitochondria). Combined with the abnormal distribution of mitochondria at metaphase, this occasionally led to extreme outcomes, such as QL.pp inheriting either no mitochondria or much more than usual (**Fig. S7c**). Additionally, the morphological difference between QL.pa and QL.pp mitochondria with respect to sphericity (PC3) is lost, with only a significant difference in volume (PC1) remaining (**Fig. 2e**).

We observed that the loss of *drp-1* also dysregulates mitochondrial segregation albeit in a different way. Average mitochondrial density is significantly lower in QL.pp compared to QL.p and QL.pa (0.026 vs 0.032 and 0.033) (**Fig. 2c**, *drp-1*(*bc455*), resulting in a significant increase in average mitochondrial density ratio between metaphase and post-cytokinesis (from 1.85 to 2.31) (**Fig. 2d**). However, compared to wild type, the differences in mitochondrial densities and the increase in mitochondrial density ratio observed in *drp-1*(lf) animals are smaller. As in *fzo-1*(lf), the difference in sphericity of mitochondria (PC3) between the two daughter cells is lost, with only a significant difference in volume (PC1) remaining (**Fig. 2e**). In contrast to *fzo-1*(lf), there is no correlation between mitochondrial density ratios at metaphase and post-cytokinesis (**Fig. S3**). Time series of QL.p divisions reveal that mitochondria in QL.p are often hyper-fused into one large mitochondrion that encircles the aligned chromosomes at metaphase (**Fig. S8a-c**). When QL.p divides, QL.pp sometimes inherits a large fraction of this hyper-fused mitochondrion, whereas other times it receives only a small fraction. Consequently, mitochondrial segregation during QL.p division is not predictable.

Together, these results indicate that mitochondrial fission and to a lesser extent mitochondrial fusion contribute to the equal mitochondrial distribution in QL.p at metaphase and the unequal mitochondrial segregation during QL.p division.

To confirm these findings, we analysed mitochondrial distribution in QL.p at metaphase and in QL.p daughter cells post-cytokinesis in *fzo-1*(lf) and *drp-1*(lf) animals using microfluidic-based immobilization. As shown in **Fig.S9b** and **Fig.S10b**, the phenotypes observed were essentially identical to those observed using nanobead-based immobilization (**Fig. 2**). For example, in *fzo-1*(lf) animals, no significant increase in average mitochondrial density ratio was observed between metaphase and post-cytokinesis (1.89 and 1.79) and in *drp-1*(lf) animals, compared to wild type, a significant but smaller increase was observed (increase from 1.08 to 1.48 in *drp-1*(lf) compared to an increase from 0.94 to 3.02 in *+/+*).

Using microfluidics-based immobilization also allowed us to determine whether the loss of mitochondrial dynamics affects the equal segregation of mitochondria during QL.pa division. As in wild type, QL.pa division in both *fzo-1*(lf) and *drp-1*(lf) animals is essentially symmetric by cell volume (**Fig.S9c** top left, bottom right; **Fig. S10c**, top left, bottom right). In addition, we found that in both mutants, mitochondria are segregated essentially equally as indicated by similar average mitochondrial densities in QL.pa and its daughters QL.paa and QL.pap, as well as similar average mitochondrial density ratios before and after QL.pa division (**Fig. S9c**, top centre and right; **Fig. S10c**, top centre and right). As observed in wild type, we also found significant correlations between mitochondrial density ratios at metaphase and post-cytokinesis in both *fzo-1*(lf) and *drp-1*(lf) animals (**Fig. S9c**, bottom left; **Fig.S10**, bottom left). Therefore, equal mitochondrial segregation during QL.pa division is unaffected by the loss of either mitochondrial fission or fusion.

### Unequal mitochondrial segregation during QL.p division is not determined by mitochondrial membrane potential

To determine whether there is a difference in mitochondrial membrane potential (Δψ) between mitochondria inherited by QL.pa and QL.pp, we used staining with tetramethylrhodamine ethyl ester (TMRE, orange) and a transgene that labels the cell membrane (myristoylated SFmTurquoise2ox, magenta), chromatin (SFmTurquoise2ox-HistoneH1, magenta) and mitochondria (mtGFP, cyan) (*bcIs158* transgene) (**Fig. 3a**). TMRE staining was combined with nanobead-based immobilization and 3D rendering of cell and mitochondrial volumes to measure mitochondrial membrane potential (i.e. TMRE fluorescence intensity) within individual organelles (see Methods) (**Fig. 3a**).

**Figure 3.**
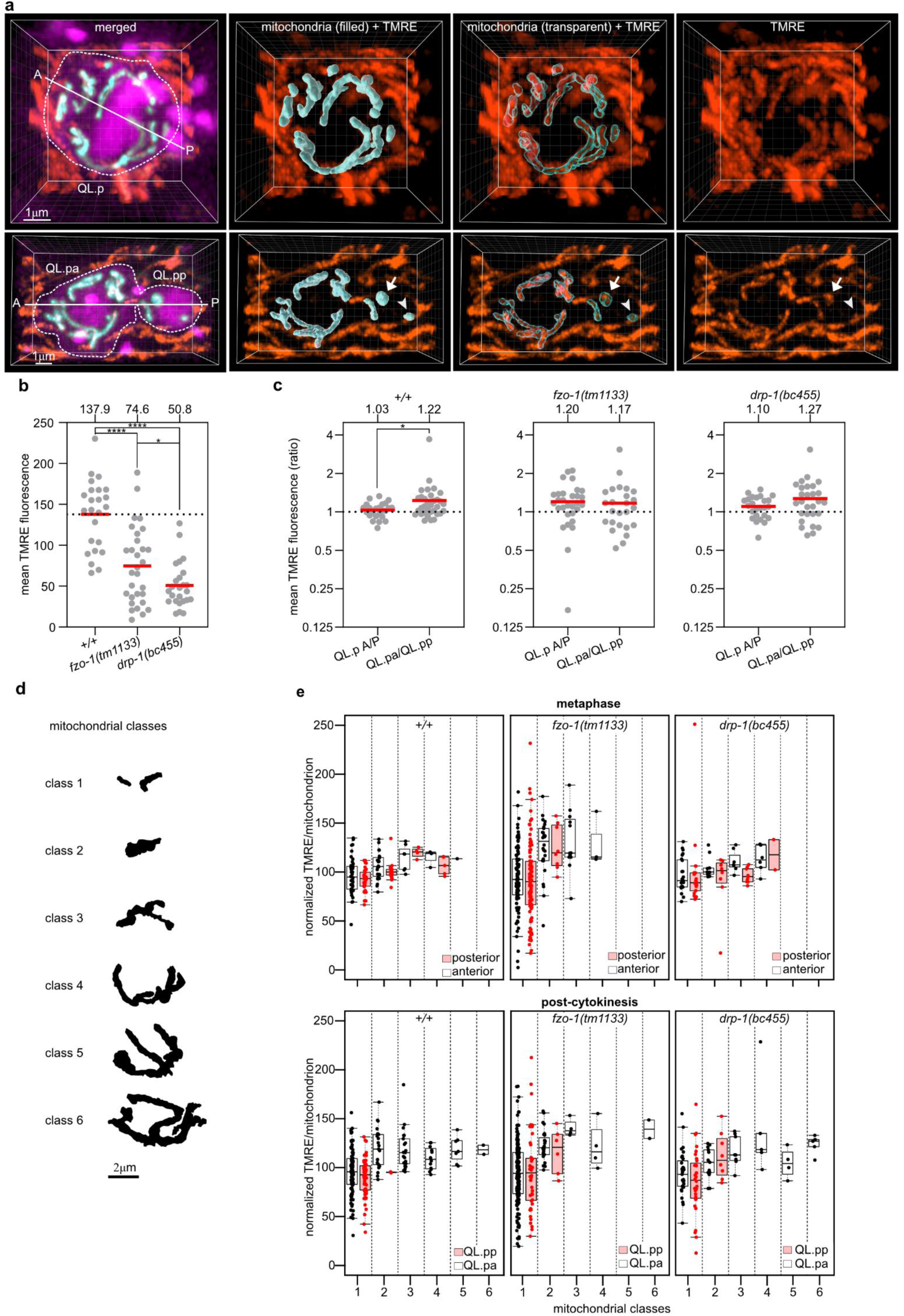
(previous page). Mitochondrial membrane potential of partitioned mitochondria is random during QL.p division. **a**) Representative 3D image of wild-type QL.p labelled with TMRE at metaphase (top row) and post-cytokinesis (bottom row). In both top and bottom left images (merged): both myristoylated SFmTurquoise2ox (cell membrane) and SFmTurquoise2ox::his-24 (chromatin) are shown in magenta, mtGFP (mitochondria) in cyan, and TMRE in orange (the same colouring is used in Figure S11a,b, and c). The A-P axes show the orientation of QL.p division. Arrows and arrowheads point to mitochondria with higher and lower TMRE intensities, respectively. TMRE fluorescence intensities are measured within each volume of mitochondria (see “mitochondria (3D) + TMRE” images) in Imaris. **b**) Comparison of the average TMRE fluorescence intensity in wild-type and mutant (*fzo-1(tm1133)* and *drp-1(bc455)*) animals. **c**) Comparison of the mean TMRE fluorescence intensity ratio between metaphase (QL.p A/P) and post-cytokinesis (QL.pa/QL.pp) in wildtype (+/+), fzo-1(tm1133), and drp-1(bc455). **d**) Outline of the six mitochondrial volume classes used in Figure 3e. The outlines refer to the examples given in figure S11. **e**) Box and whisker plots of normalized TMRE fluorescence intensity per mitochondrion at metaphase (top plots) and post-cytokinesis (bottom plots) in wildtype (left plots), fzo-1(tm1133) (centre plots), and drp-1(bc455) (right plots). Mitochondria were sorted into six volume classes to investigate the difference in mitochondrial activity between organelles with different volume. The mean TMRE intensity within each mitochondrion was normalized by dividing it with the mean TMRE intensity of all mitochondria in the respective cell (QL.p for metaphase or QL.pa and QL.pp, together, for post-cytokinesis) (see methods for further information). P values are calculated using the Kruskal-Wallis test with the Benjamini, Krieger and Yekutieli correction (b), and the Mann-Whitney test (c). Normality was tested with the Shapiro-Wilk test. *: P value ≤ 0.05; **: P value ≤ 0.01; ***: P value ≤ 0.001; ****: P value ≤ 0.0001. In panels b and c, red lines and dots represent the population average and the individual samples’ average, respectively. In panel e, dots represent the individual mitochondria. All data are from animals expressing *bcIs158* transgene.

We determined the average TMRE fluorescence intensities of mitochondria in the anterior and posterior halves of QL.p at metaphase in wild type and found that they are similar (QL.p A/P ratio = 1.03). However, post-cytokinesis, mitochondria in QL.pa showed a slightly but significantly higher TMRE fluorescence intensity than mitochondria in QL.pp (QL.pa/QL.pp ratio = 1.22) (**Fig. 3c**, +/+). The difference in average TMRE fluorescence intensities observed between QL.pa and QL.pp appears to suggest that mitochondria are selectively segregated based on membrane potential. However, we found that TMRE fluorescence intensity positively correlates with mitochondrial volume and negatively correlates with sphericity (**Fig. S11a,b**). Therefore, to correct for size- and shape-related effects, we binned individual mitochondria into six volumetric classes, where class 1 represents small and spherical mitochondria and class 6 large and tubular mitochondria (see Methods) (**Fig.3d**, **Fig.S12a-c**). Importantly, when we compared mitochondria of the same volumetric class, we did not find statistically significant differences in TMRE fluorescence intensities between posterior and anterior mitochondria in QL.p at metaphase or between mitochondria in QL.pa and QL.pp post-cytokinesis (**Fig. 3e**, metaphase, post-cytokinesis, *+/+*; note, each data point refers to one mitochondrion). This suggests that the difference in average TMRE fluorescence intensities observed between QL.pa and QL.pp is the result of morphological differences (rather than differences in membrane potential) between mitochondria in QL.pa and QL.pp. This also suggests that unequal mitochondrial segregation during QL.p division is not determined by mitochondrial membrane potential.

We also investigated the effect of altered mitochondrial dynamics on mitochondrial membrane potential during the division of QL.p (see **Fig.S13**). Average TMRE fluorescence intensities (as measured in QL.p cells at metaphase) are significantly lower in *fzo-1(tm1133)* and *drp-1(bc455)* mutants than in wild type (**Fig. 3b**, *fzo-1*(*tm1133*), *drp-1*(*bc455*)). Furthermore, we found that the difference between metaphase and post-cytokinesis with respect to mean TMRE fluorescence ratio observed in wild type is lost in *fzo-1*(lf) and *drp-1*(lf) mutants (**Fig. 3c**). In both mutants, the distribution of TMRE intensities among different classes of mitochondria at metaphase are overall similar to those in wild type with a broader range of intensities observed in *fzo-1*(lf) animals (**Fig. 3e**, metaphase). After division, as in wild type, class 1 mitochondria with high and low TMRE intensities are inherited by both QL.p daughters in *fzo-1*(lf) and *drp-1*(lf) mutants (**Fig.3e,** post-cytokinesis). However, both mutants are characterized by a higher frequency of class 2 mitochondria in QL.pp and relatively fewer class 3-6 mitochondria in QL.pa (**Fig. 3e**, post-cytokinesis). These changes in the frequencies of mitochondrial classes after division likely explain the loss of asymmetry in average TMRE fluorescence ratio between QL.p daughters in these mutants.

Overall, these results suggest that the difference in TMRE fluorescence intensities detected between QL.pa and QL.pp is not the result of biased mitochondrial segregation based on mitochondrial membrane potential during QL.p division. Instead, the difference in membrane potential between QL.pa and QL.pp is probably determined by the difference in mitochondrial morphologies, which are dependent on mitochondrial fusion and fission. Consequently, disrupting mitochondrial dynamics impacts mitochondrial segregation not only in terms of density and morphology, but also membrane potential.

### Unequal mitochondrial segregation during QL.p division correlates with the acquisition of the cell death fate by QL.pp

Since QL.p distributes mitochondria unequally between the two daughter cells and the two daughter cells acquire distinct cell fates, we investigated whether unequal mitochondrial segregation correlates with the divergence of these fates. For this experiment, we used microfluidic-based immobilization for long-term tracking of Q lineage cells and acquired quantitative data on (i) QL.p cell cycle length (time between QL division and QL.p division), (ii) QL.pa cell cycle length (time between QL.p division and QL.pa division) as a measure of QL.pa fate, (iii) QL.pp survival time (time between QL.p division and QL.pp death; see **Fig. S14** and Methods for how QL.pp survival time was determined) as a measure of QL.pp fate, (iv) mitochondrial segregation during QL.p division and (v) mitochondrial segregation during QL.pa division (see **Fig. 4a**). QL.p and its daughter cells were tracked using an ultrafast confocal mode. Super-resolution imaging during QL.p and QL.pa division was performed at metaphase and post-cytokinesis (Methods).

**Figure 4.**
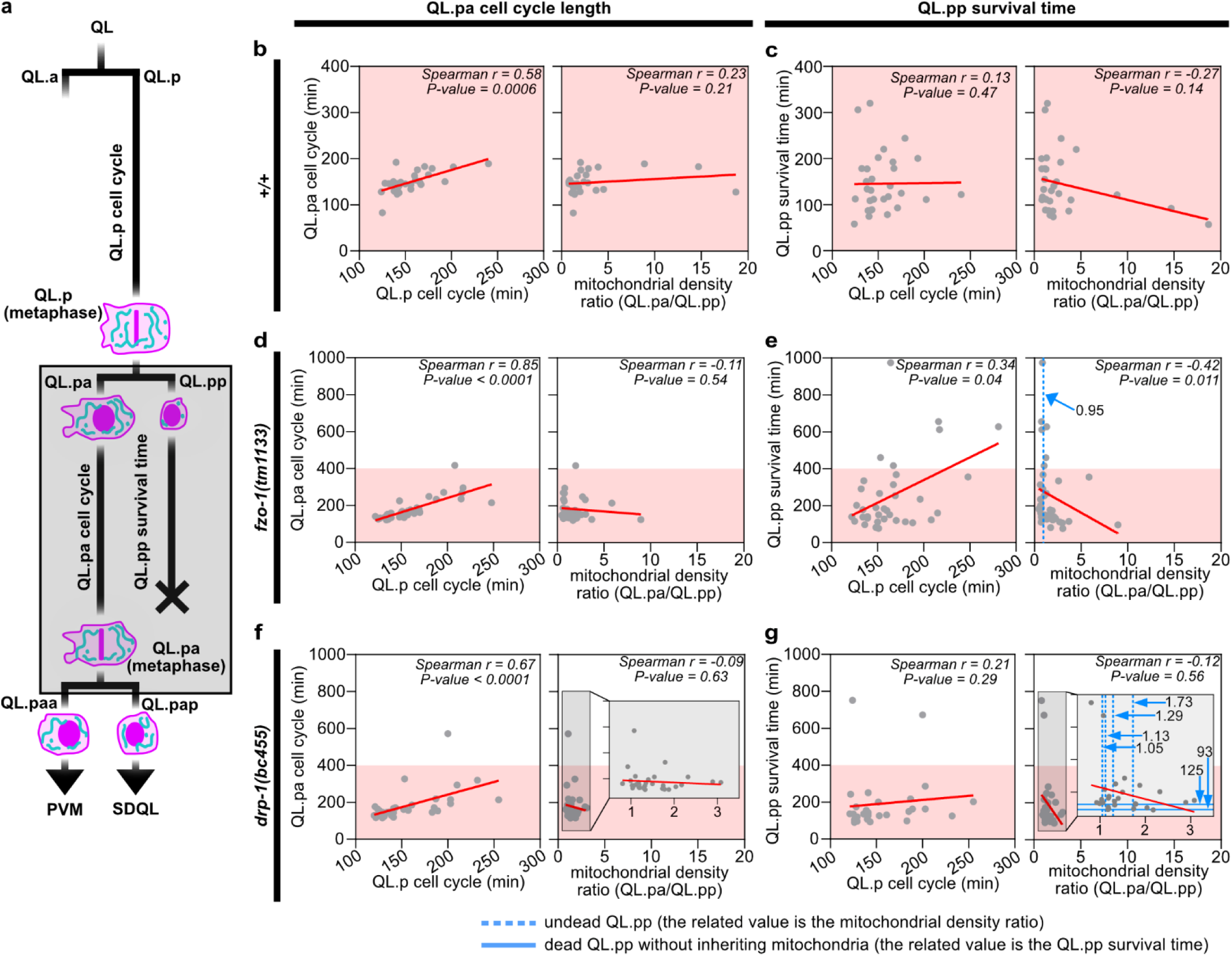
Analysis of the relationship between mitochondrial partitioning and QL.pa and QL.pp fate. **a**) Schematics of the QL.p lineage recorded in wild-type (*+/+*), *fzo-1(tm1133)* and *drp-1(bc455)* animals expressing *bcIs153* transgene. The grey box highlights QL.pa and QL.pp cell fates, which were followed in super resolution (metaphase and post-cytokinesis time points for QL.pa) and in high resolution (morphological change (=death) for QL.pp (see Fig. S10)), respectively. QL.p division was also followed in super resolution at metaphase and post-cytokinesis. **b,c**) correlation between QL.pa cell cycle length (**b**) or QL.pp survival time (**c**) and QL.p cell cycle length (left) or mitochondrial density ratio (right) in wildtype (*+/+*). **d,e**) correlation between QL.pa cell cycle length (b) or QL.pp survival time (c) and QL.p cell cycle length (left) or mitochondrial density ratio (right) in *fzo-1(tm1133)*. **f,g**) correlation between QL.pa cell cycle length (**b**) or QL.pp survival time (**c**) and QL.p cell cycle length (left) or mitochondrial density ratio (right) in *drp-1(bc455)*. Red lines = linear regression fitted on scattered plots. Dots represent data from single QL.p divisions. All correlations were analysed calculating the Spearman correlation coefficient *r* and its related p-value. n = 31, 35, and 33 in wild-type (*+/+*), *fzo-1(tm1133)* and *drp-1(bc455)*, respectively. Data were collected from animals expressing *bcIs153* transgene.

We found that in wild-type animals, the cell cycle lengths of QL.pa and QL.p strongly correlate (**Fig. 4b**), while QL.pp survival time does not correlate with QL.p cell cycle length (**Fig. 4c**). Furthermore, average mitochondrial density ratio does not correlate with QL.pa cell cycle length (**Fig.4b**), yet it shows a negative but non-significant correlation with QL.pp survival time (**Fig. 4c**). This suggests that QL.p and QL.pa cell cycles may progress at a rate proportionate to the overall developmental rate of the animal. In contrast, QL.pp survival time is independent of developmental rate, but may negatively correlate with the level of asymmetry in mitochondrial segregation. In other words, a smaller mitochondrial density ratio (i.e. QL.pp inherits relatively more mitochondria) may correlate with longer QL.pp survival times. Conversely, a larger mitochondrial density ratio (i.e. QL.pp inherits relatively fewer mitochondria) may correlate with shorter QL.pp survival times.

Next, we analysed *fzo-1(tm1133)* and *drp-1(bc455)* animals, in which unequal mitochondrial segregation is compromised and more variable. As in wild type, QL.pa cell cycle lengths strongly correlates with QL.p cell cycle length in both *fzo-1*(lf) and *drp-1*(lf) animals, but mitochondrial density ratio does not correlate with QL.pa cell cycle length (**Fig. 4d,f**). In *fzo-1*(lf), the survival time of QL.pp shows a weak positive correlation with QL.p cell cycle length (**Fig. 4e**), but this is not the case for *drp-1*(lf) (**Fig. 4g**). In addition, in *fzo-1*(lf), average mitochondrial density ratio shows a significant negative correlation with QL.p survival time **(Fig. 4e**). In *drp-1*(lf), as in wild type, average mitochondrial density ratio shows a negative but non-significant correlation with QL.pp survival time (**Fig.4g**). Importantly, in both *fzo-1*(lf) and *drp-1*(lf) mutants, we captured QL.p lineages in which QL.pp inappropriately survived. In the *fzo-1*(lf) background, we captured one QL.p lineage in which QL.pp survived. This particular QL.pp cell inherited relatively more mitochondria with an average QL.pa/QL.pp mitochondrial density ratio of 0.95 rather than the average 1.74 (**Fig. 4e**, dotted vertical blue line). In the *drp-1*(lf) background, we captured four QL.p lineages in which QL.pp survived. Three of these QL.pp cells inherited relatively more mitochondria with average QL.pa/QL.pp mitochondrial density ratios between 1.05 and 1.29 rather than the average 1.48 (**Fig. 4g**, dotted vertical blue lines). (The fourth QL.pp cell inherited slightly fewer mitochondria with an average QL.pa/QL.pp mitochondrial density ratio of 1.78.) In the *drp-1*(lf) background, we also captured two lineages, in which QL.pp did not inherit any mitochondria. In both cases, QL.pp died markedly faster with a survival time of only 93 min or 125 min, compared to an average survival time of 208 min in *drp-1*(lf) (**Fig. 4g**, horizontal blue lines and **Fig. S15**, left).

To be able to include the cases of inappropriately surviving QL.pp cells in *fzo-1(tm1133)* and *drp-1(bc455)* animals, we conducted a survival analysis. A multivariate Cox proportional-hazards regression^24^ was used to investigate the relationship between QL.pp survival time and both QL.pp mitochondrial density and genotype. We found that there was a significant effect of QL.pp mitochondrial density on its survival time as indicated by relative death rate (**Fig. 5a**). Increasing QL.pp mitochondrial density from 2.8% (average for wild-type) to 7.1% is predicted to halve its relative death rate (from 1.0 to 0.5) (**Fig. 5a**). QL.pp survival probabilities of the different genotypes are shown in the Kaplan-Meier survival probability plot (**Fig. 5b**). The loss of *fzo-1* or *drp-1* shows a significant effect on QL.pp survival probability compared to wild type (*+/+*) (**Fig. 5b**; p-value=0.0044). In wild-type animals, essentially all QL.pp cells die within 300 min post-cytokinesis. In contrast, in *fzo-1*(*tm1133*), some QL.pp cells die only after 900 min, and in *drp-1*(*tm1108*), some survive longer than 1200 min.

**Figure 5.**
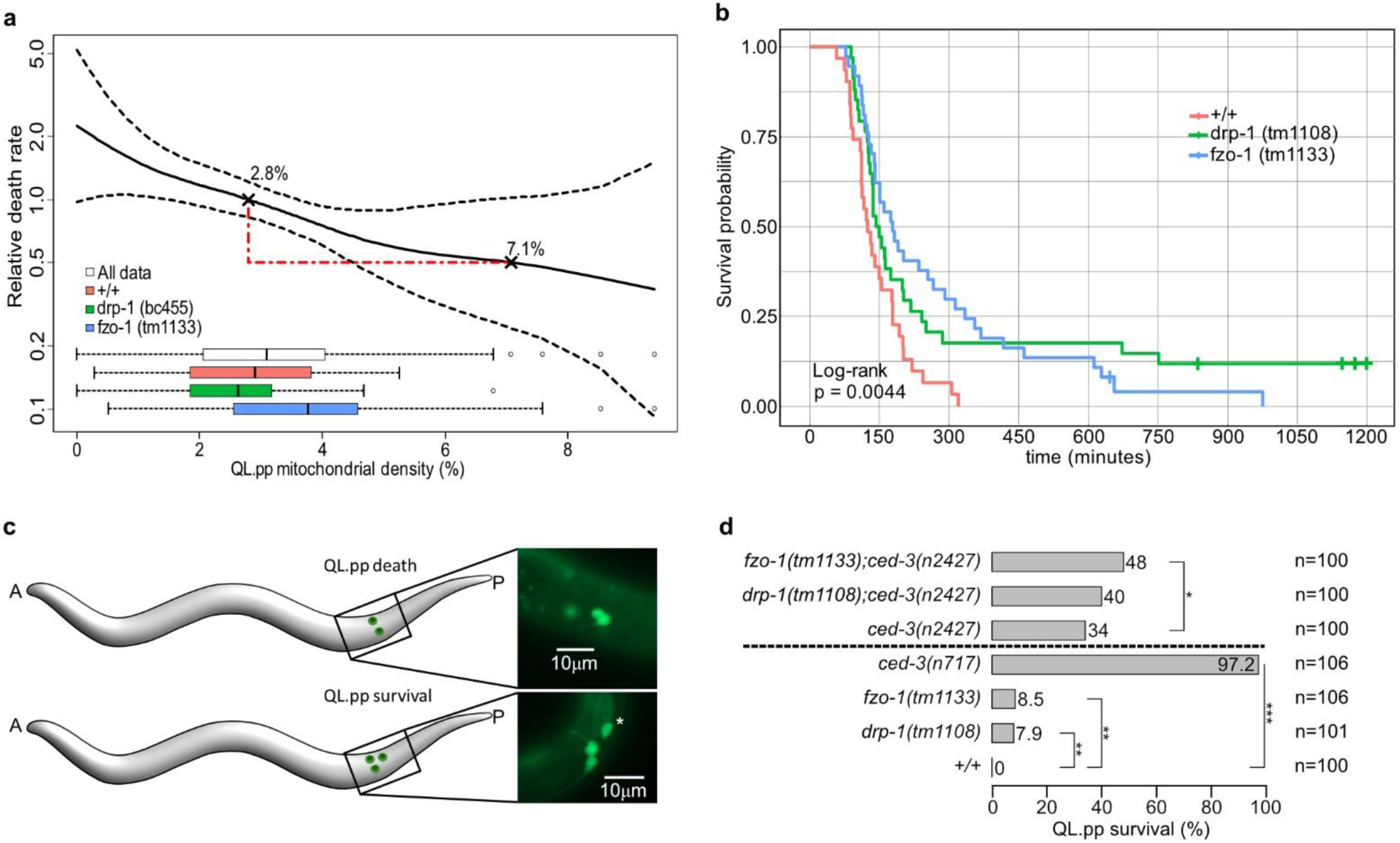
Impact of mitochondrial density and mitochondrial dynamics on QL.pp cell death fate. **a**) Relative death rate explainable by the mitochondrial density in QL.pp, as obtained by the Cox proportional hazards regression. Dotted lines indicate pointwise standard errors. The spread of the data across different genotypes is illustrated by the inset boxplots. Death rates are relative to the average mitochondrial density in QL.pp (2.8%; Figure 1b), with the example showing a halving of death rate in QL.pp at a mitochondrial density of 7.1%. **b**) Kaplan-Meier survival curves for wild-type (*+/+*), *fzo-1(tm1133)*, and *drp-1(bc455)*. Ticks along the curves represent QL.pp cells that were phenotypically determined to survive and subsequently no longer followed (right censored data). Global log rank test indicates significant differences in survival between the genotypes. **c**) Schematics of the QL.pp survival assay. Cell counting is conducted in early L2 larvae, were only two cells (top) are visible (QL.paa (PVM) and QL.pap (SDQL)) in wild-type animals. Whenever QL.pp survives, a third cells is counted (asterisk) (bottom) or even four cells can be counted (divided QL.pp, example not shown). **d**) QL.pp survival comparisons between wildtype (*+/+*), *drp-1(tm1108)*, *fzo-1(tm1133)* and *ced-3(n717)* (lower side) and *fzo-1(tm1133);ced-3(n2427)*, *drp-1(tm1108);ced-3(n2427)* and *ced-3(n2427)* (upper side). P values refer to Fisher’s Exact tests:. *: P value ≤ 0.05; **: P value ≤ 0.01; ***: P value ≤ 0.001; ****: P value ≤ 0.0001. Data in Figure 5a,b and Figure 5c,d derive from *bcIs153*- and *bcIs133*-expressing animals, respectively. In panels a and b, n = 31, 36, and 37 in wild-type (*+/+*), *fzo-1(tm1133)* and *drp-1(bc455)*, respectively.

In summary, these analyses in *drp-1*(lf) and *fzo-1*(lf) mutants reinforce the notion that inheriting relatively more mitochondria correlates with longer QL.pp survival times and inheriting relatively fewer mitochondria with shorter QL.pp survival times.

Interestingly, when accounting for both genotype and mitochondrial density simultaneously using the multivariate Cox regression, the loss of *drp-1* shows a significant effect on survival (Cox proportional hazard: −0.7571 ± 0.2705 SE, p = 0.005). However, the loss of *fzo-1* no longer shows a significant effect on survival once mitochondrial density is accounted for (−0.4889 ± 0.2651 SE, p = 0.065). This suggests that the effect of the loss of *fzo-1* on QL.pp survival is exclusively due to *fzo-1*’s contribution to mitochondrial segregation during QL.p division and, hence, QL.pp mitochondrial density. Conversely, the effect of the loss of *drp-1* on QL.pp survival is not only due to *drp-1*’s contribution to QL.pp mitochondrial density but a second mitochondrial density-independent feature of QL.p division and/or QL.pp.

Finally, we verified the findings obtained from the micro-fluidic experiment and the analyses of relative death rate and survival probability of QL.pp by conducting a QL.pp survival assay using transgenic animals that express a Q lineage-specific GFP transgene^25^ (*bcIs133* transgene) **(Fig. 5c)**. In wild type, QL.pp always dies (0% inappropriate survival) (**Fig.5d**, +/+), whereas in control animals lacking the caspase *ced-3* (*ced-3(n717)*), 97.2% of QL.pp cells inappropriate survive (**Fig. 5d**). We found that both *fzo-1(tm1133)* and *drp-1(tm1108)* animals exhibited low, but significant, inappropriate QL.pp survival compared to control animals (8.5 and 7.9%, respectively). In the background of the weak *ced-3* loss-of-function mutation *n2427*, neither *fzo-1(tm1133)* nor *drp-1(tm1108)* showed a synergistic effect in terms of inappropriate QL.pp survival (**Fig. 5d**). Overall, animals defective for mitochondrial dynamics show significant inappropriate QL.pp survival (**Fig. 5b**).

In conclusion, our findings are consistent with the notion that unequal mitochondrial segregation during QL.p division and, hence mitochondrial density in QL.pp, correlates with the acquisition of the cell death fate by QL.pp. Specifically, low mitochondrial density in QL.pp is associated with very rapid QL.pp death, whereas high density coincides with inappropriate QL.pp survival.

## Discussion

### Mitochondria segregate unequally during asymmetric QL.p division

Unequal mitochondrial segregation has been demonstrated in budding yeast but has been difficult to study in animal systems. Taking advantage of the highly reproducible development of *C. elegans* and advances in live-cell super-resolution microscopy and immobilization methods, we have developed methodology that enables us to quantitatively study mitochondrial segregation in real time and at single organelle resolution during asymmetric cell divisions in *C. elegans* larvae. Using this methodology, we have obtained highly reproducible data, even when using different transgenes, immobilization methods, growth media and bacterial foods. These data demonstrate that mitochondrial segregation during the asymmetric division of the neuroblast QL.p is unequal by density and morphology. Consequently, compared to its sister cell (QL.pa), QL.pp has a lower density of mitochondria and its mitochondria are predominantly small and spherical. To the best of our knowledge, this is the first *in vivo* demonstration of unequal mitochondrial segregation during asymmetric cell division in an animal system. We also detected lower membrane potential per mitochondrial volume in QL.pp; however, our data suggests that this is due to the overrepresentation of small and spherical mitochondria in QL.pp.

We previously showed that cells programmed to die during *C. elegans* development are characterized by ‘fragmented’ (i.e. small and spherical) mitochondria^14^. Ding Xue and colleagues proposed that this is the result of the general dismantling of cells undergoing caspase-dependent apoptotic cell death^27^. Furthermore, they reported that mitochondrial fission is the result of CED-3 caspase-dependent cleavage and activation of DRP-1 in apoptotic cells. We now present evidence that the ‘fission’ of mitochondria in a cell programmed to die, QL.pp, does not occur post-cytokinesis. Instead, it is the result of unequal mitochondrial segregation by density and morphology during the asymmetric cell division that gives rise to QL.pp.

### Mitochondrial segregation during asymmetric QL.p division is non-random

We propose that mitochondrial segregation during QL.p division is an actively controlled process and therefore non-random. First, we found that during symmetric QL.pa division, mitochondrial distribution in the daughter cells (QL.paa and QL.pap) is predictable based on mitochondrial distribution in QL.pa at metaphase. This suggests that during QL.pa division, mitochondria segregate as preconfigured by their distribution at metaphase. Since mitochondria are equally distributed throughout the anterior and posterior halves of QL.pa, mitochondria are equally segregated into QL.paa and QL.pap. In contrast, during asymmetric QL.p division, mitochondrial distribution in the daughter cells (QL.pa and QL.pp) is not predictable based on mitochondrial distribution in QL.p at metaphase. Since mitochondria are also equally distributed throughout the anterior and posterior halves of QL.p, mechanisms must be at work that control mitochondrial segregation and cause unequal mitochondrial distribution between QL.pa and QL.pp. Indeed, we present evidence that these mechanisms include anteriorly directed mitochondrial transport and localized mitochondrial fission (see below, **Fig.6**). Second, the loss of mitochondrial fusion (i.e. *fzo-1*(*tm1133*) background) makes mitochondrial distribution in QL.pa and QL.pp predictable based on mitochondrial distribution in QL.p. Importantly, daughter cell size asymmetry is maintained in *fzo-1*(*tm1133*) mutants (**Fig. S9**). Therefore, the processes that establish daughter cell size asymmetry are not sufficient to cause unequal mitochondrial distribution in QL.pa and QL.pp, at least in the *fzo-1*(*tm1133*) background. This argues against a model in which unequal mitochondrial segregation is caused by daughter cell size asymmetry. Finally, mitochondrial distribution in QL.pa and QL.pp also becomes predictable in *pig-1*(*gm344*) animals, and this may at least in part be due to a loss of anteriorly directed mitochondrial transport and localized mitochondrial fission. The PAR-1-like kinase PIG-1 MELK is required for QL.p polarization along the anterior-posterior axis and the establishment of daughter cell size asymmetry during QL.p division. We found that anteriorly directed mitochondrial transport, localized mitochondrial fission and the establishment of daughter cell size asymmetry occur simultaneously during QL.p division (see **Fig.2a** and **Fig.S4**). Therefore, we consider it likely that anteriorly directed mitochondrial transport and localized mitochondrial fission are lost in *pig-1*(lf) mutants as a result of the loss of *pig-1*’s role in QL.p polarization, rather than its role in the establishment of daughter cell size asymmetry. Interestingly, *Xenopus* MELK has been shown to localize to mitochondria^26^. It is, therefore, conceivable that PIG-1 may also play a direct role in mitochondrial segregation, for example, in anteriorly directed mitochondrial transport.

**Figure 6.**
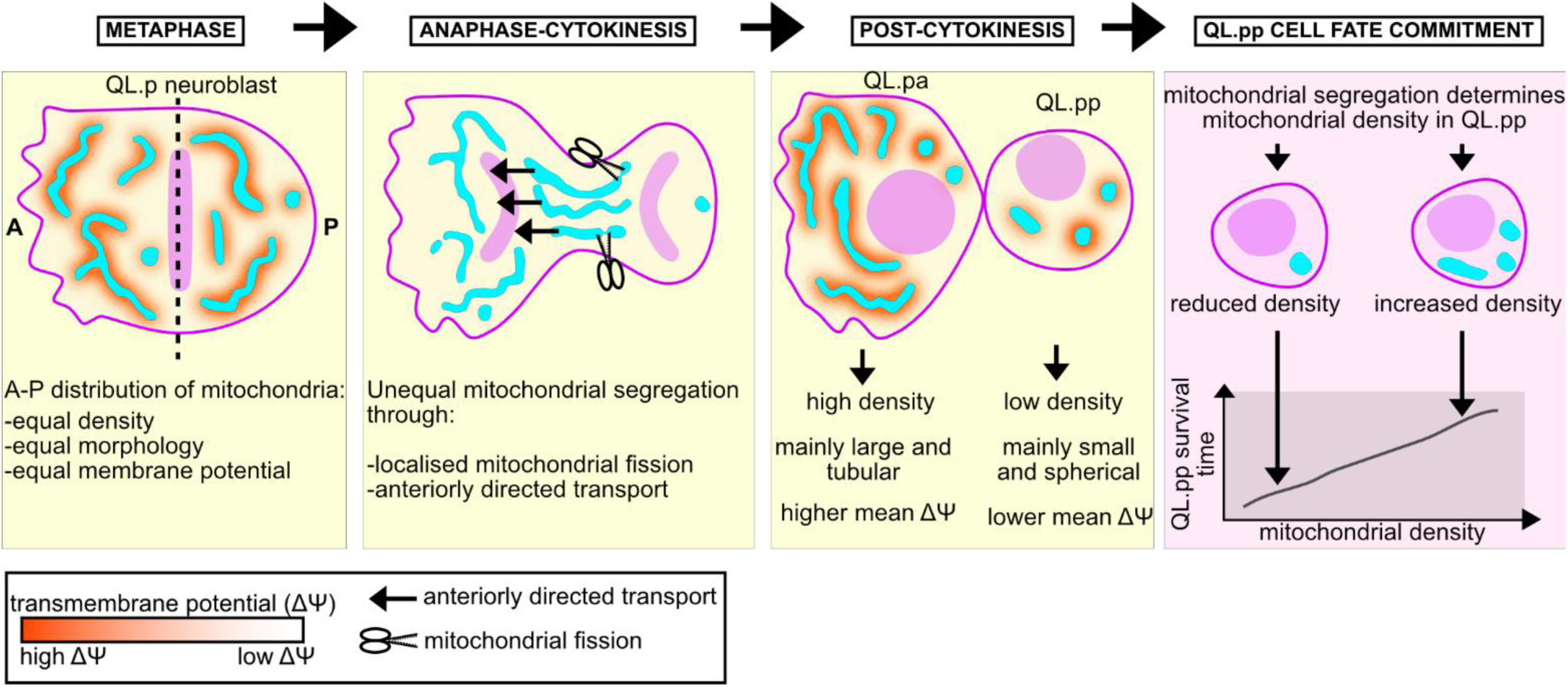
Working model of mitochondrial segregation during QL.p division. Mitochondria are symmetrically distributed at metaphase in terms of density, morphology, and membrane potential. Mitochondrial dynamics regulates mitochondrial distribution at metaphase and the asymmetric activity of mitochondria fission regulates asymmetric mitochondrial segregation during QL.p division. The asymmetric inheritance of mitochondria may be determined by selective (active) mechanisms that transport and/or tether mitochondria into the surviving daughter cells. On the contrary, segregation appears stochastic for membrane potential. As a result, the larger surviving daughter cell inherits more mitochondria than the smaller fated-to-die daughter cell. The higher mitochondrial density in the larger daughter is mainly determined by the enrichment of large tubular mitochondria that are characterized on average by higher membrane potential. On the other hand, the smaller daughter cell is left with relatively fewer mitochondria, the quantity of which can partially determine if and how quickly QL.pp acquires the cell death fate.

We propose the following model for how mitochondria are segregated during asymmetric QL.p division (**Fig.6**). At metaphase, mitochondria are equally distributed throughout the anterior and posterior halves of QL.p with respect to density, morphology and membrane potential. This equal distribution is dependent on the ability of mitochondria to divide and fuse and, hence, most likely on ‘normal’ mitochondrial morphology. During QL.p division, small and spherical mitochondria become enriched in the posterior half, which will form the smaller daughter QL.pp; this is dependent on at least two processes, both of which are dependent on PIG-1 MELK: (i) mitochondrial fission in the posterior half and (ii) anteriorly directed transport of larger and more tubular mitochondria. What could activate mitochondrial fission in the posterior half of QL.p as it divides? We previously showed that a low level of CED-3 caspase is activated in QL.p and that prior to QL.p division, an anteroposterior gradient of this CED-3 caspase activity is established in QL.p with higher levels of CED-3 caspase in the posterior half of QL.p^25^. Furthermore, as stated above, Ding Xue and colleagues reported that DRP-1 can be activated through CED-3 caspase-dependent cleavage^27^. Hence, mitochondrial fission could be activated in the posterior half of QL.p through localized CED-3-dependent DRP-1 cleavage. Concerning anteriorly directed transport of mitochondria, we propose that it is mediated by cytoskeletal elements and specific motor proteins. Our results also suggest that small and spherical mitochondria on the one hand and hyper-fused mitochondria on the other are not competent for anteriorly directed transport. Therefore, we propose that anteriorly directed transport is selective based on mitochondrial morphology. Finally, because we observed no difference in mitochondrial membrane potential when comparing mitochondria of similar volumes and sphericities in QL.pa and QL.pp post-cytokinesis, our results do not support the notion that transport is selective by mitochondrial membrane potential.

### Unequal mitochondrial segregation during asymmetric QL.p division contributes to cell fate divergence in daughter cells

As a result of unequal mitochondrial segregation during QL.p division, compared to its sister cell (QL.pa), QL.pp has a lower density of mitochondria. We propose that this lower density of mitochondria in QL.pp is required for the timely and reproducible death of QL.pp and, hence, the divergence of the QL.pa and QL.pp fates during asymmetric QL.p division. Using three different genetic backgrounds (wild type, *fzo-1*(lf), *drp-1*(lf)), we consistently found that changes in mitochondrial density in QL.pp impact the ability of QL.pp to adopt the cell death fate. Specifically, we found that decreased mitochondrial density in QL.pp correlates with QL.pp dying faster and, conversely, that increased mitochondrial density in QL.pp correlates with QL.pp dying slower, or not at all. It remains to be determined whether changes in mitochondrial density are sufficient to cause changes in QL.pp survival time. However, our findings are consistent with the idea that mitochondrial density impacts the activation of the apoptosis pathway in QL.pp and/or the execution of apoptotic cell death in QL.pp. Of note, the accompanying changes in mitochondrial densities in QL.pa had no impact on QL.pa cell cycle time. However, as a result of QL.pa’s size (∼3 times the size of QL.pp), these changes may have been too subtle to have an impact. Consistent with this, in wild-type animals (microfluidics-based immobilization), mitochondrial densities in QL.pa range from 0.033 to 0.068 units and in QL.pp from 0.003 to 0.053 units.

Cell to cell variability in mitochondrial volume (rather than mitochondrial density) has previously been proposed to contribute to cellular noise and differences for example in transcriptional rates in mammalian cells^30,32^. Furthermore, mitochondrial volume or density has also been proposed to impact the activation of apoptosis in mammalian cells. However, while one study found that mitochondrial density correlates with resistance to TRAIL-induced apoptosis^34^ (which would be consistent with our findings in *C. elegans*), another study found that cells with increased mitochondrial volume are more prone to die^36^. One possible reason for the discrepancy between these studies could be the fact that cellular volume was not taken account in the second study.

Why might increasing mitochondrial density in QL.pp cause QL.pp to die slower or not at all? The anti-apoptotic Bcl-2 protein CED-9 localises to the outer mitochondrial membrane^38^. Therefore, CED-9 levels are expected to increase in QL.pp with increasing mitochondrial density. Furthermore, CED-9 is a substrate of CED-3 caspase^40^, which we have confirmed *in vitro*^42^. Therefore, it is conceivable that increased CED-9 levels in QL.pp compete with other CED-3 caspase substrates, thereby compromising the ability of CED-3 caspase to trigger the execution of apoptotic cell death. Alternatively, apoptotic cell death might be compromised by other mitochondrial proteins or mitochondrial metabolites, which are also expected to increase in QL.pp with increasing mitochondrial density. Changes in levels of anti-apoptotic CED-9, other mitochondrial proteins or mitochondrial metabolites may also explain why QL.pp dies faster when mitochondrial density is decreased or – in some cases – no mitochondria are inherited by QL.pp. How the apoptosis pathway is fully activated in QL.pp in the absence of mitochondria is an enigma, unless it is triggered and the CED-4 Apaf1 apoptosome assembled prior to the completion of QL.p division. (We consider it unlikely that the low level of CED-3 caspase activity that is specifically inherited by QL.pp as a result of the formation of a CED-3 caspase gradient in QL.p [see above] is sufficient to kill QL.pp, even less so to kill QL.pp faster than at normal mitochondrial density.)

Our data also suggest that in contrast to *fzo-1*, *drp-1*’s impact on QL.pp survival may not be limited to its role in mitochondrial segregation during QL.p division. We previously showed that DRP-1 (but not FZO-1) can interact with CED-9 Bcl-2 when bound by the BH3-only protein EGL-1 and that this interaction promotes DRP-1-dependent mitochondrial fission^44^. Therefore, we posit that CED-9/EGL-1- and DRP-1-dependent mitochondrial fission may contribute to the cell death fate of QL.pp independently of DRP-1’s contribution to unequal mitochondrial segregation. Alternatively, the CED-9/EGL-1/DRP-1 complex may contribute to unequal mitochondrial segregation in a fission-independent manner.

In conclusion, we provide the first evidence that the non-random segregation of mitochondria during asymmetric cell division can contribute to cell fate divergence in a developing animal. Hence, the non-random segregation of mitochondria represents a novel intrinsic mechanism for cell fate divergence in the context of asymmetric cell division.

## Acknowledgments

We thank Alex Hajnal (University of Zurich, Switzerland) and Andrew deMello (ETH Zurich, Switzerland) for their support of S.B.

## Methods

### Strains and genetics

Unless noted otherwise, all *C. elegans* strains were cultured at 20°C as described^28^. Bristol N2 was used as the wild-type strain. Mutations and transgenes used in this study are: LGII: *fzo-1(tm1133)*^29^. LGIV*: ced-3(n2427)*^31^, *ced-4(n717)*^33^, *drp-1(bc455)* (this study), *drp-1(tm1108)*^29^, *pig-1(gm344)*^21^, *bcIs153* (DOI https://doi.org/10.21203/rs.3.rs-4320882/v1) (P_toe-2::_mtGFP::unc-54 3’UTR (DOI https://doi.org/10.21203/rs.3.rs-4320882/v1) + P_egl-17_::Myri-mCherry::pie-1 3’UTR^50^ + P_egl-17_::mCherry-TEV-S::his-24^50^ + rol-6(su1006)^35^), *bcIs158* (this study) [P_toe-2::_mtGFP::unc-54 3’UTR (DOI https://doi.org/10.21203/rs.3.rs-4320882/v1) + P_egl-17_::Myri-SFmTurquoise2ox::pie-1 3’UTR (DOI https://doi.org/10.21203/rs.3.rs-4320882/v1) + P_egl-17_::SFmTurquoise2ox-TEV-S::his-24 (DOI https://doi.org/10.21203/rs.3.rs-4320882/v1) + rol-6(su1006)^35^], and *bcIs133*^25^. Additional transgenes used in this study are: *bcIs159* (this study) [P_mab-5_::EYFP-CUP-5::unc-54 3’UTR (this study) + P_egl-17_::Myri-SFmTurquoise2ox::pie-1 3’UTR + P_egl-17_:: SFmTurquoise2ox -TEV-S::his-24 + rol-6(su1006)^35^], *bcIs160* (this study) [P_toe-2::_mtGFP::unc-54 3’UTR + P_toe-2_::SP-12mCherryKDEL::unc-54 3’UTR (this study) + P_egl-17_::Myri-SFmTurquoise2ox::pie-1 3’UTR + P_egl-17_::SFmTurquoise2ox-TEV-S::his-24 + rol-6(su1006)^35^] and *bcIs162* (this study) [P_toe-2_::TOMM-20(N-term)-mKate2::tbb-2 3’UTR (DOI https://doi.org/10.21203/rs.3.rs-4320882/v1) + P_egl-17_::Myri-SFmTurquoise2ox::pie-1 3’UTR + P_egl-17_:: SFmTurquoise2ox -TEV-S::his-24 + rol-6(su1006)^35^]. Throughout our studies, we used information and tools available on WormBase (https://wormbase.org/#012-34-5)^52,54^.

### Plasmid construction

All plasmids, inserts and oligos used are listed below (end of Methods section). Amplicons were generated through Q5 PCR (CDS) or LongAmp PCR (promoter sequences). **pBC1978** (P_mab-5_::EYFP::cup-5::unc-54 3’UTR): the plasmid pBC1977 (P_mab-5_::unc-54-3’UTR) was digested with NheI and BstEII to generate the vector backbone. The EYFP insert was amplified from pBC1689 template with oligos IS-EYFP-N-term-F and IS-EYFP-N-term-R. The cup-5 insert was amplified from pBC1983 (P_mab-5_::cup-5::EYFP::unc-54 3’UTR) with oligos CUP-5-for-Nterm-EYFP-F and CUP-5-for-Nterm-EYFP-R. The two inserts were inserted into the vector backbone through Gibson Assembly. The cup-5 CDS was inserted into pBC1983 by removing its 5^th^ intron (2659bp), between exons 5 and 6, containing the uncharacterized gene *R13A5.15*. **pBC1984** (P_toe-2_::SP12-mCherry-KDEL::unc-54 3’UTR): the plasmid pBC1590 was digested with NheI and NotI to generate the vector backbone. The SP12-mCherry-KDEL insert was amplified from pBC986 template with oligos NheI-For and NotI-Rev and then digested with NheI and NotI. The insert was inserted into the vector backbone through T4 ligation.

### CRISPR-Cas9-mediated generation of the *bc455* allele

In order to CRISPR-Cas9 the *drp-1(tm1108)* deletion, we adapted a recently published methodology^37^. All RNA molecules are listed below (end of Methods section). The *tm1108* deletion variant was generated by selecting two crRNAs named ISsgRNA1(tm1108) and ISsgRNA2(tm1108) that were identified using CRISPOR 5.01 (input sequence ce11-chrIV-5538489-5540480, 1991bp long, PAM: NGG). The repair template IStm1108REPAIRDNA was designed such that both “arms” of the ssDNA had 40-50bp overlap (lower case) with both blunt ends after cutting out the deletion sequence, while the central part (upper case) consists of a small duplication of the 5’ end that naturally occurred during the original TMP/UV mutagenesis of *drp-1*^29^. The protocol consisted of three main steps prior to microinjection. Preparation of MIX1 (incubation at 95°C for 1 minute + cooling): 50μM 5’ crRNA (ISsgRNA1(tm1108)) (2μl), 50μM 5’ crRNA (ISgsRNA2(tm1108)) (2μl) and 100μM tracrRNA (2μl). Preparation of MIX2 (incubation at 37°C for 15 minute): MIX1 (1.5μl), Cas9 (62μM) (0.4μl) and dpy-10 crRNA (50μM) (1μl). Preparation of MIX3, which was spun down at 12000g for 3 minutes and the top 7μl were transferred into a new tube as the final microinjection solution: MIX2 (2.9μl), IStm1108REPAIRDNA (1μl) and 1x TE, pH 7.5 (IDT) (5.5μl). The MIX2 contains the *dpy-10* crRNA as a coinjection marker to select those animals (dumpy phenotype) in which CRISPR-Cas9 gene editing was successful^39^. Animals were outcrossed once with N2 (wild-type) animals to remove the *dpy-10* mutation in homozygosity (*bcIs158*) or self-crossed to remove the *dpy-10* mutation in heterozygosity (*bcIs153*). crRNAs were synthesized by Merck with the addition of a 5’ modification (2’-O-methyl and phosphorothioate linkages) for extra stability and purified by HPLC. crRNAs were ordered without premixing with tracrRNA. Cas9 and tracrRNA were synthesised by IDT.

### Mechanical immobilization of L1 larvae

The immobilization and imaging protocols used to study QL.p division in wild type, *pig-1(gm344)*, *fzo-1(tm1133)* and *drp-1(bc455)* in super-resolution and at high temporal resolution (**Fig. 2** and related figures) are the same as reported in *Segos et al.* (DOI https://doi.org/10.21203/rs.3.rs-4320882/v1). For *bcIs160* and *bcIs159; bcIs162* animals (**Fig. S1**), the immobilization protocol is the same as reported in *Segos et al.* (DOI https://doi.org/10.21203/rs.3.rs-4320882/v1).

### Image processing and 3D rendering from recordings acquired on animals mechanically immobilized

The image processing protocol is the same as reported in *Segos et al*. (DOI https://doi.org/10.21203/rs.3.rs-4320882/v1) but performing 5 iterations of the Richardson-Lucy deconvolution algorithm instead of 20 iterations. We observed that reducing the number of iterations reduced the risk of artificial fission of mitochondrial volumes during 3D rendering in Imaris.

### Confocal microscopy and 2D image analysis (mitochondria + ER experiment)

We used an upright LSM980 with an Airyscan2 setup with a GaAsP-TMP detector and performed Super Resolution Airyscan with a C Plan Apochromat 63x/1.40 objective. mCherry, GFP and SFmTorquoise2ox fluorophores were excited respectively through 594nm (0.5%), 488nm (0.2%), and 405nm (0.5%) wavelengths, while their detection ranges were always 300-720nm. We adjusted other settings as follows: FOV = 13.47×13.47µm (382×382pixels), voxel size = 0.035×0.035×0.130µm, pixel time = 0.66µs (frame time = 449.4ms), detector gain = 850V, and scan direction = bidirectional (no averaging). Z-stacks were Airyscan-processed (2D, standard strength) before export. We did not perform a pairwise acquisition of metaphase and post-cytokinesis for the ER images, due to suspected phototoxicity of SP12-mCherry-KDEL and mtGFP resulting in the appearance of circular mitochondria during post-cytokinesis (images not shown). Images were then processed in Fiji (ImageJ v1.53f51) through background subtraction (rolling ball radius = 60pixels). We drew ROIs along cell boundary on each slice of the z-stack and we measured area and Integrated Density within each ROI and summed them up (for all slices relative to the QL.p anterior side, QL.p posterior side, QL.pa and QL.pp). We then calculated the mean Integrated Density (Integrated Density / area) from summed areas and Integrated Densities to obtain the overall density of fluorescence intensities (a proxy for organelle density) at both metaphase and post-cytokinesis. The summed areas were used to calculate respective cell size at both metaphase and post-cytokinesis. We used summed areas and densities of fluorescence intensities to measure organelle ratios and cell volume ratios at metaphase and post-cytokinesis.

### Validation of the lysosome-specific marker

We validated the lysosome-specificity of the EYFP::CUP-5 marker by co-labelling lysosomes with lysotracker Red DND-99® (**Fig.S2**). Not all lysotracker-labelled lysosomes are visible with EYFP::CUP-5 because transcription of this reporter gene is driven by the promoter region of *mab-5*. This gene encodes a homeobox protein (TF) that controls the formation of the posterior-specific pattern of cells during the post-embryonic development of *C. elegans*^41^. *mab-5* is expressed only in a subset of cells found in the posterior half of L1 larvae, among which QL descendants^43,45^. We calculated the Manders colocalization coefficient (MCC) of EYFP::CUP-5 fluorescence with that of lysotracker Red. On average, the unthresholded MCC was 0.91 (sd ±0.02), while the thresholded MCC was 0.81 (sd±0.16), which is similar to the MCC previously obtained for cemOrange::cup-5^46^.

### Confocal microscopy and 2D image analysis (mitochondria + lysosome experiment)

We used an upright LSM980 with an Airyscan2 setup and a GaAsP-TMP detector and performed Super Resolution Airyscan with a C Plan Apochromat 63x/1.40 objective. mKate2, EYFP and SFmTorquoise2ox fluorophores were excited respectively through 594nm (0.4%), 514nm (0.7%), and 445nm (0.7%) wavelengths, while their detection ranges were always 300-720nm. We adjusted other settings as follows: FOV = 12.54×12.54µm (324×324pixels), voxel size = 0.039×0.039×0.130µm, pixel time = 0.72µs (frame time = 409.59ms), detector gain = 850V, and scan direction = bidirectional (no averaging). Z-stacks were Airyscan-processed (2D, standard strength) before export. We performed a pairwise acquisition of metaphase and post-cytokinesis images, which were then processed in Fiji through background subtraction (rolling ball radius = 60pixels). We drew ROIs along cell boundary on each slice of the z-stack and we measured area and Integrated Density within each ROI and summed them up (for all slices relative to the QL.p anterior side, QL.p posterior side, QL.pa and QL.pp). We then calculated the mean Integrated Density (Integrated Density / area) from summed areas and Integrated Densities to obtain the overall density of fluorescence intensities (as proxy of organelle density) at both metaphase and post-cytokinesis. The summed areas were used to calculate respective cell size at both metaphase and post-cytokinesis. We used summed areas and density of fluorescence intensities to measure organelle ratios and cell volume ratios at metaphase and post-cytokinesis.

### TMRE staining of L1 larvae

NGM plates with TMRE were prepared one day prior to imaging. NGM agar was melted and TMRE (InvitrogenTM T669) was added to a final concentration of 10nM. NGM+TMRE was then poured into small petri plates (±1/2ml) and left to cool for 2-3 hours in the dark. Next, *E. coli* OP50 was suspended in M9 and pipetted on plates (20μl). L1s animals expressing *bcIs158* were synchronized for 1hr by washing out all adults and letting eggs hatch for 1hr on medium NGM plates. Freshly hatched L1s were washed off the plates with MPEG and pipetted on bacterial lawns grown overnight on NGM+TMRE plates. After 2.5 hours of growth at 20°C, L1s were washed off and transferred on a seeded medium NGM plate without TMRE for another 2.5 hours (or longer for slow growing strains) at 20°C. Mounting and immobilization were performed following the protocol outlined in Segos et al (DOI https://doi.org/10.21203/rs.3.rs-4320882/v1).

### Confocal microscopy and 3D image analysis (TMRE experiment)

We used an upright LSM980 with an Airyscan2 setup with a GaAsP-TMP detector and performed Super Resolution Airyscan with a C Plan Apochromat 63x/1.40 objective. TMRE, GFP, and SFmTorquoise2ox fluorophores were excited at 561nm (1.0%), 488nm (0.9%), and 405nm (0.8%) respectively, while their detection ranges were always 300-720nm. We adjusted other settings as follows: FOV = 12.62×12.62µm (358×358pixels), voxel size = 0.035×0.035×0.130µm, pixel time = 0.66µs (frame time = 449.4ms), detector gain = 850V, and scan direction = bidirectional (no averaging). Z-stacks were Airyscan-processed (2D, standard strength) before export. We did not perform a pairwise acquisition of metaphase and post-cytokinesis images due to suspected phototoxicity of TMRE and mtGFP resulting in the occasional appearance of circular mitochondria post-cytokinesis (images not shown). Images of the mtGFP channel in Fiji only were processed through two consecutive background subtractions (rolling ball radius = 50pixels). Average TMRE fluorescence values were calculated per cell from actual TMRE fluorescence within mitochondrial volumes in Imaris and were used to compare different genetic backgrounds. Since L1 larvae absorbed the TMRE dye at different levels due to inter-animal variability in feeding and metabolic activity, mitochondria of QL.p in distinct animals are not comparable across sample, without normalization. Therefore, comparisons between mitochondria within individual cells were performed by normalizing TMRE fluorescence (=TMRE fluorescence that is relative to the respective TMRE average in QL.p at metaphase or at post-cytokinesis). The normalization was done by measuring the average TMRE fluorescence per cell (overall average of QL.p anterior and QL.p posterior sides (metaphase) or overall average of QL.pa and QL.pp combined (post-cytokinesis)) and the normalized TMRE fluorescence was calculated as: *(individual TMRE fluorescence/average TMRE fluorescence)*100*. Next, the normalized TMRE fluorescence was paired with the respective mitochondrion volume and sphericity. Finally, we classified mitochondria by splitting the min-max volume range of the control data sets (metaphase and post-cytokinesis) into 6 morphological classes (class 1: round (=fragmented) mitochondria; class 6: tubular/networked (=fused) mitochondria). The same volume intervals were applied to mutant data sets.

### Microfluidics: L1 larvae preparation

For all experiments, we adapted the methodology described in *Berger et al. (2021)*^20^ for use with L1 larvae.

#### L1 larvae preparation

L1 larvae were prepared by washing out all adults and larvae from medium plates (4x or more plates) with MPEG (5.8 g Na2HPO4, 3g KH2PO4, 0.5g NaCl, 1g NH4Cl to 1l MilliQ water and autoclave, then add 1g Polyethylene glycol 8000 (MPEG 0.1%)). We let embryos hatch for 90 minutes and collected L1s by washing plates with MPEG (2ml for 4-5 plates by transferring buffer+L1s plate-by-plate) into a 1.5ml microcentrifuge tube(s). We next filtered L1s through a 10μm (pluriStrainer Mini 10 μm, PluriSelect) and pipetted the filtered MPEG+L1s into 1.5ml tubes. We then centrifuged animals at 400g for 3min and discarded the MPEG leaving L1s in the pellet (∼20μl). We next added 1.7ml of S-Basal and centrifuged at 400g for 3min. Subsequently, we discarded the S-Basal leaving 50/60µl with pelleted L1s, which were used for microfluidics mounting.

#### Food preparation

We added 0.2μm-filtered LB in a 50ml Falcon tube to 20ml final volume (2x falcons = 40ml). Next, we added 45μl of *E. coli* NA22 culture (OD_600_ = 1.9; kept at 4°C) to both Falcon tubes and let them grow for 16.5hrs in a 37°C shaking incubator (final OD_600_ was always between 1.7-2.0). The OD_600_ was measured using a spectrophotometer (Thermo Scientific Genesis 30 Visible). Bacteria were spun down at 3000g (or rcf) (15.3rad) for 5min (Eppendorf Centrifuge 5804R). We discarded the liquid and resuspended the pellet in 2ml of S-Basal by pipetting up and down and then combined the two 2ml volumes. We spun down the tube at 3000g for 3min, discarded the liquid and resuspend the pellet in 4ml of S-Basal. We repeated this step once more. We then resuspend the pellet in 1ml S-Basal and transferred this into a 2ml microcentrifuge tube. We added OptiPrep (0.65ml) and S-Basal+PluronicF127 (1%w/v) (0.33ml) to the bacterial suspension and mixed by gently pipetting up and down. (Fast pipetting can make the bacterial preparation toxic for L1 animals.) We next filtered the *E. coli* NA22 solution twice, once through a 10 μm filter and next through a 5 μm filter (pluriStrainer Mini 10 μm or 5 μm, PluriSelect).

#### Microfluidics device operation

We mounted L1 larvae in the L1 chip as described before^20^ and followed their development throughout the experiment (up to 22 hours). We manually switched between Multiplex Airyscan imaging (see below) and Airyscan Super Resolution (see next two sections) whenever QL.p and QL.pa were at metaphase and at post-cytokinesis. To achieve super-resolution image quality, we manually actuated the on-chip compression layer (20psi and 15psi during QL.p and QL.pa divisions, respectively) prior to switching to super-resolution mode. We did not compress the system at any other time (i.e.: while following the cell in Multiplex Airyscan mode) to minimize mechanical stress. We took images every 3.5 minutes. We modified the MATLAB coding of the Arduino MICRO (ATmega32U4 hacking) microcontroller and connected it to an “ON/OFF” external button to manually control the system.

### Characterization of QL.pp cell death confocal microscopy images

Using the adapted methodology originally described in *Berger et al. (2021)*^20^, we imaged wild-type animals expressing the *bcIs153* transgene (Fig. S14a,b). No compression was applied during the entire recording, and images were taken every 5.5 minutes for 12 hours total. We used an inverted LSM980 with an Airyscan2 setup with a GaAsP-TMP detector (LSM) and Multialkali-PMT detector (DIC). We performed LSM + DIC (Nomarsky) with a C Plan Apochromat 63x/1.40 objective. We set the High Contrast (DIC III) Wollastone prism for DIC and manually set all microscope components (aperture, condenser, polarizer and Wollastone prism) to optimize DIC image quality. LSM mode was selected for acquisition of the mCherry signal (561nm excitation wavelength, laser power = 0.08%), while DIC images were acquired through a transmission channel (T-PMT) using the same wavelength used to excite mCherry. The detection wavelengths for mCherry and the transmitted channels are 570-694nm and 300-900nm, respectively. We adjusted other settings as follows: FOV = 133.46×34.57µm (2965×768pixels), voxel size = 0.045×0.045×1.000µm, pixel time = 0.26µs (frame time = 449.74ms), detector gain = 610V, and scan direction = bidirectional (no averaging). We used DIC and LSM to follow morphological changes of QL.pp, including its transformation into a refractile cell corpse^47^. We identified a sudden morphological shift from shrunk round QL.pp to prolate spheroid corpse, which coincided with the appearance of refractility by DIC (**Fig. S14c**, white arrowheads). This morphological change was followed by the quick disappearance of the corpse (in LSM) (**Fig. S14c**, times +187min to +209min). We investigated multiplex confocal recordings and identified the exact same change (**Fig. S14d**, times +147min and +150:30min). QL.pp fluorescence follows a unimodal distribution before death (**Fig. S14e**, black profiles) and a multimodal distribution after death (**Fig. S14e**, red profiles). These fluorescence distributions coincide with the respective morphologies (**Fig. S14f**). This change in QL.pp morphology was used to determine the time of QL.pp death.

### Confocal microscopy to follow QL.pa and QL.pp fates

For this experiment, we followed the methodology described in *Berger et al. (2021)*^20^ with the changes mentioned above. We used wild-type and mutant animals expressing the *bcIs153* transgene. We used an inverted LSM980 with Airyscan2 to image wild-type and *drp-1(bc455)* animals and an upright LSM980 with Airyscan2 to image *fzo-1(tm1133)*, due to the unavailability of the inverted microscope during the experiment. We followed the entire QL.p lineage through Multiplex Airyscan SR-8Y (long-term imaging) imaging and acquired Airyscan super resolution images (see below) for metaphase and post-cytokinesis during both QL.p and QL.pa cell divisions. We performed Multiplex Airyscan with a C Plan Apochromat 63x/1.40 objective using the Q lineage markers (mCherry) only. mCherry was excited at 561nm (laser power = 0.3% (=0.03% in standard LSM mode)) and the detection wavelength was 574-720nm. The tile function was used to save multiple XYZ positions so that positions (P1, P2, P3, …) were organized and assigned across the microfluidic device geometry. We adjusted other settings as follows: FOV = 133.46×34.57µm (1992×516pixels), voxel size = 0.067×0.067×1.000µm, pixel time = 1.05µs (frame time = 159.74ms), detector gain = 990V, and scan direction = bidirectional (no averaging). After completing the experiment, recordings were 2D Airyscan-process (standard strength) in ZenBlue 3.3. Airyscan-processing was necessary to visualize and process images in ImageJ.

### Confocal microscopy to record QL.p and QL.pa divisions in super-resolution

We used an inverted or upright LSM980 with an Airyscan2 (see above) setup with a GaAsP-TMP detector and performed Super Resolution Airyscan with a C Plan Apochromat 63x/1.40 objective. mCherry and GFP fluorophores were excited at 561nm (0.6%) and 488nm (0.6%), respectively, while their detection ranges were always 300-720nm. We adjusted other settings as it follows: FOV = 20.66×7.12µm (464×160pixels), voxel size = 0.045×0.045×0.130µm, pixel time = 1.08µs (frame time = 226.89ms), detector gain = 990V, and scan direction = bidirectional (no averaging). Z-stacks were Airyscan-processed (2D, standard strength) before export. Images were processed in Fiji and 3D rendered in Imaris according to *Segos et al.* (DOI https://doi.org/10.21203/rs.3.rs-4320882/v1) with the following changes. Before channel subtraction (mtGFP channel – mCherry channel), the mCherry image was multiplied by 0.08 to reproduce the theoretical emission of mCherry (8%) when excited by the 488nm wavelength. Images were deconvolved using 10 iteration of the Richardson-Lucy algorithm.

### QL.pp survival assay

The number of surviving QL.pp cells was scored using the P_toe-2_gfp (*bcIs133*) transgene, as previously described^25^. Late L1 larvae/young L2 larvae (>10 hours post-QL.p division) were mounted on 2% agar pads using 10 mM levamisole in M9 buffer as paralytic agent. The number of GFP-positive cells was determined using a Zeiss Axioscope 2. In wild-type worms there are two GFP-positive cells representing the QL.pa daughters (PVM and SDQL neurons). Up to two extra GFP-positive cells can be seen in mutants, representing inappropriately surviving QL.pp or QL.pp daughters. To validate the counting, all GFP-positive cells were assessed in DIC to ensure they were not GFP-positive corpses. The QL.pp survival percentage represents the number of QL.pp that inappropriately survived (animals with 1 or 2 extra GFP-positive cells) divided by the sample size (number of animals analysed).

### Statistical analysis

Statistical analyses were performed using GraphPad Prism8 and R (v4.3.2). For two sample comparisons of organelle densities and organelle density ratios, student’s t-tests and Wilcoxon’s tests were used based on Shapiro-Wilk’s normality test to check the normality assumption. Pairwise t-tests and signed-rank test were used for paired data and are indicated in the figures by lines connecting paired data instead of points. Spearman’s rank correlation coefficient was used to test for the strength of monotonic relationships throughout.

For the principal component analysis, six variables of mitochondrial morphology (volume, surface area, sphericity, surface to volume ratio, and oblate and prolate ellipticity) were averaged for each cell and side of the cell (i.e. QL.p anterior, QL.p posterior, QL.pa, and QL.pp). These were then scaled to have unit variance, and principal components were then constructed with PC1 containing mostly size variation, PC2 containing predominantly variance in ellipticity, and PC3 containing predominantly variance in sphericity/surface to volume ratio. The first three principal components always contained at least 92% of all variation for all genotypes studied, the remaining principal components were not considered for analysis. To test for differences following this exploratory analysis, linear models were constructed with antero-posterior location (or QL.pa-QL.pp) as a factor, residuals were inspected visually, and analysis of variance (ANOVA) was carried out.

For the QL.pp survival assay (cell counts), one-sided Fisher’s exact tests were used to compare relevant genotypes and false discovery rate was used to correct for multiple testing. For the survival analysis of the microfluidics imaging, time until death was recorded. As the five surviving cells could not be followed indefinitely, and the data was considered right censored as consequence. Cox proportional hazard’s models were fitted using the *survival* package^48,49^, and goodness of fits were explored graphically using the *survminer* package^51^. The Schoenfield and Martingale residuals were explored visually, as well as explicitly tested for. The eventual Cox regression model correlates survival time in minutes, by genotype and with mitochondrial density as a covariate using smoothing splines with three degrees of freedom. The latter was done to satisfy the linearity assumption of covariates in the cox proportional hazards model. This term, the mitochondrial density spline, is visualised as the relative death risk by mitochondrial density in Fig.5a.

### Figures and illustrations

Figures and schematics were generated in Affinity Designer 1.7.3, GraphPad Prism8 and R 4.3.2, and the Kaplan-Meier curves were plotted using the ggplot2 package^53^.

### List of oligos and RNA molecules used for cloning and mutagenesis in this study

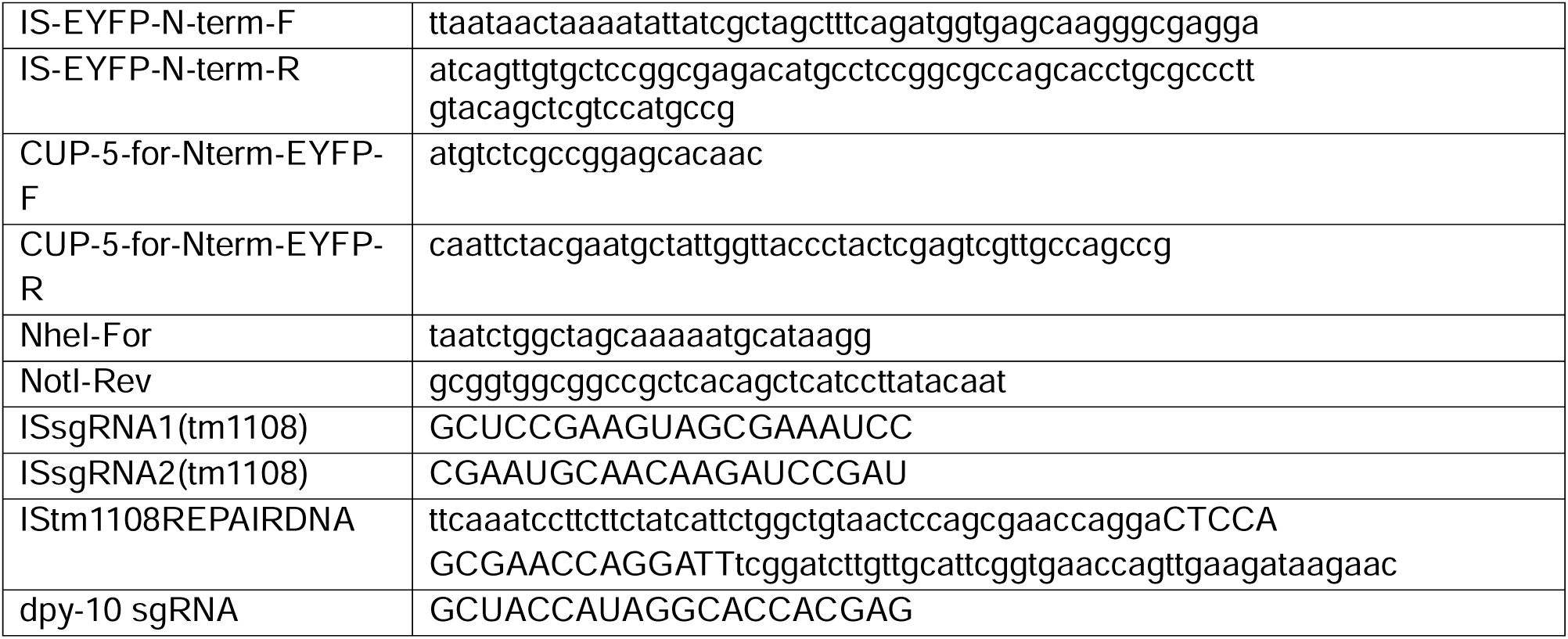

---

**Figure S1.**
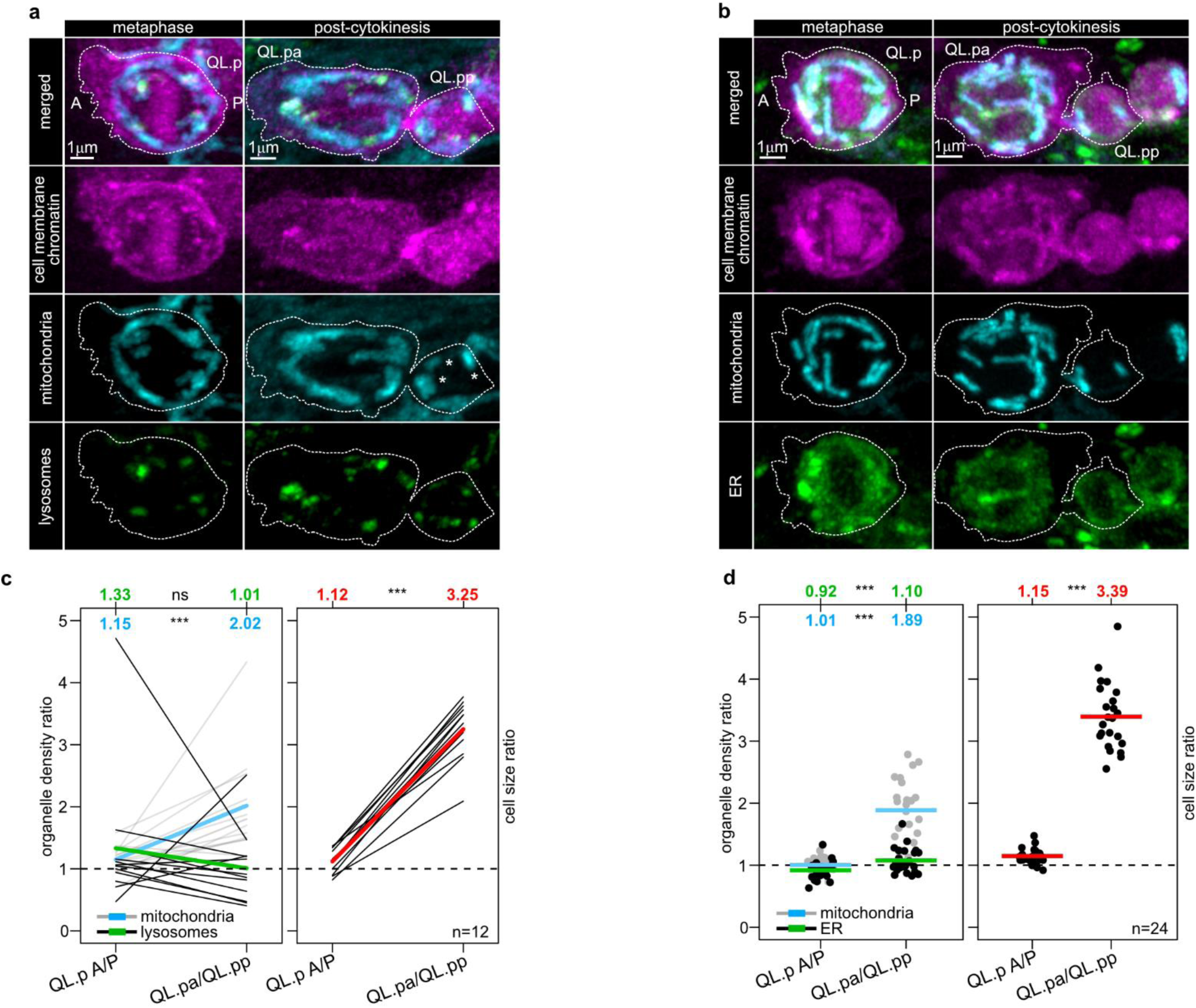
Partitioning of lysosomes and ER during QL.p division. **a**) Representative images of QL.p at metaphase and post-cytokinesis with chromatin (SFmTurquoise2ox::his-24) and cell membrane (myristoylated SFmTurquoise2ox) labelled in magenta, mitochondria (tomm-20::mKate2) in cyan, and lysosomes (eyfp::cup-5) in green (see main text for further information). **b**) Representative images of QL.p at metaphase and post-cytokinesis with chromatin (SFmTurquoise2ox::his-24) and cell membrane (myristoylated SFmTurquoise2ox) labelled in magenta, mitochondria in cyan (mtGFP), and ER (SP12-mCherry-KDEL) (in green (see main text for further information). **c,d**) Analysis of organelles density ratio at metaphase and post-cytokinesis (left), and cell size ratio (right). Measurements of organelles and cell size are based on the mean integrated density and the area (µm^2^) on each section of z-stacks, respectively, and not on volume. P values are calculated using the Wilcoxon matched pairs signed rank test (**c**, left), the paired t-test (**c**, right), the unpaired t-test (**d**, left (mitochondria), and **d**, right), and the Mann-Whitney test (**d**, left (ER)). Normality was tested with the Shapiro-Wilk test. *: P value ≤ 0.05; **: P value ≤ 0.01; ***: P value ≤ 0.001; ****: P value ≤ 0.0001. Red, cyan and green lines (and respective numbers) = average (**c,d**). Black and grey dots and lines represent data from individual QL.p divisions (**c,d)**. Data in Figure S1a,c derives from animals expressing *bcIs159* and *bcIs162* alleles, while data in Figure 5b,d derive from *bcIs160*-expressing animals.

**Figure S2.**
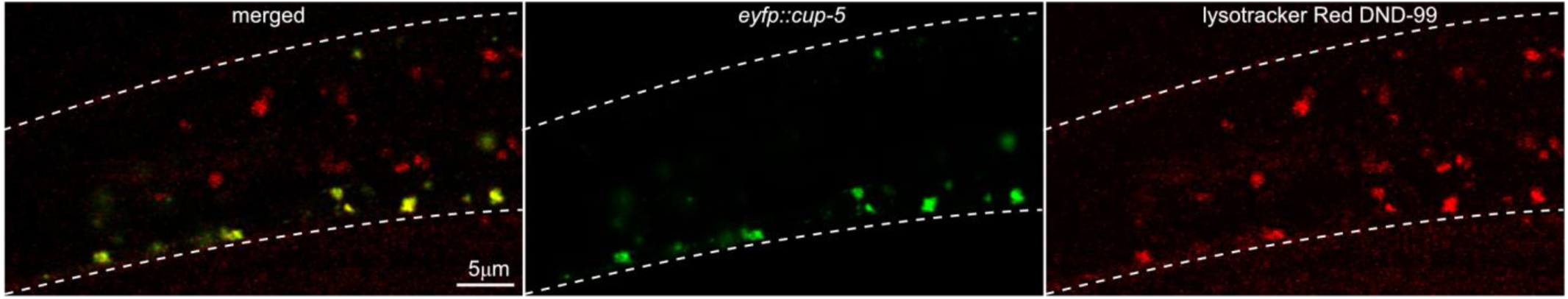
*eyfp::cup-5* transgene labels specifically lysosomes. Colocalization of *eyfp::cup-5* reporter expression with lysotracker Red DND-99 in the posterior side of an L1 larva expressing *bcIs159* transgene. This *eyfp::cup-5* transgene labels lysosomes in some cells, but not all, because its expression is driven by *mab-5* promoter, which is active only in those respective cells, among which posterior Q neuroblasts.

**Figure S3.**
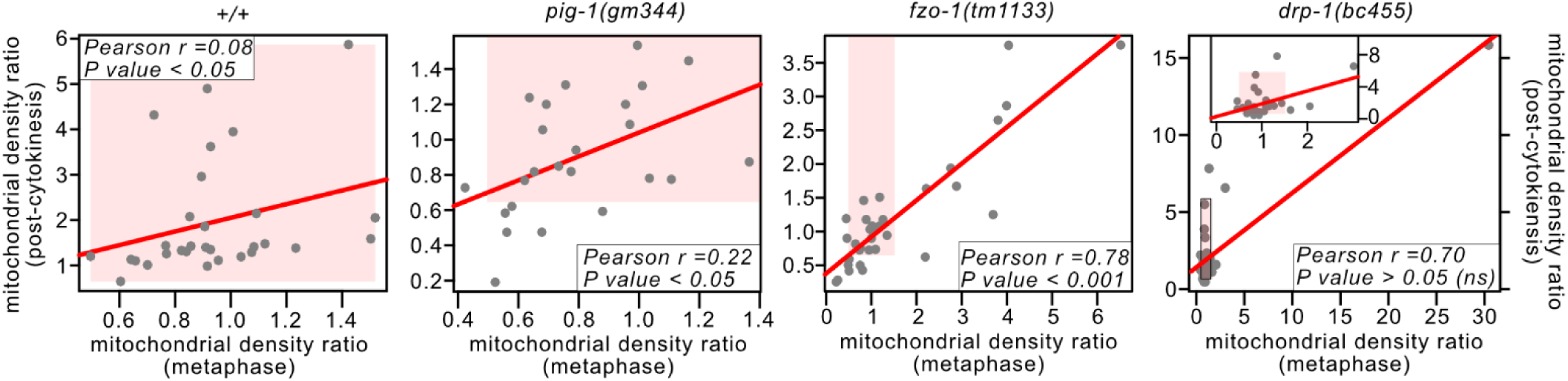
Correlation between QL.p mitochondrial density ratios at metaphase (QL.pa A/P) and post-cytokinesis (QL.pa/QL.pp). Red lines = linear regression fitted to scatter plots (dots represent values from individual QL.p divisions). Pearson’s or Spearman’s tests for association between samples were performed where appropriate, indicated by the coefficient of determination (r2) or the square of Spearman’s rho (ρ2) are respectively given, illustrating the proportion of shared variance. Normality was tested using the Shapiro-Wilk test, and homoscedasticity was tested using Bartlett’s test. In all genotypes, animals express the *bcIs153* transgene.

**Figure S4.**
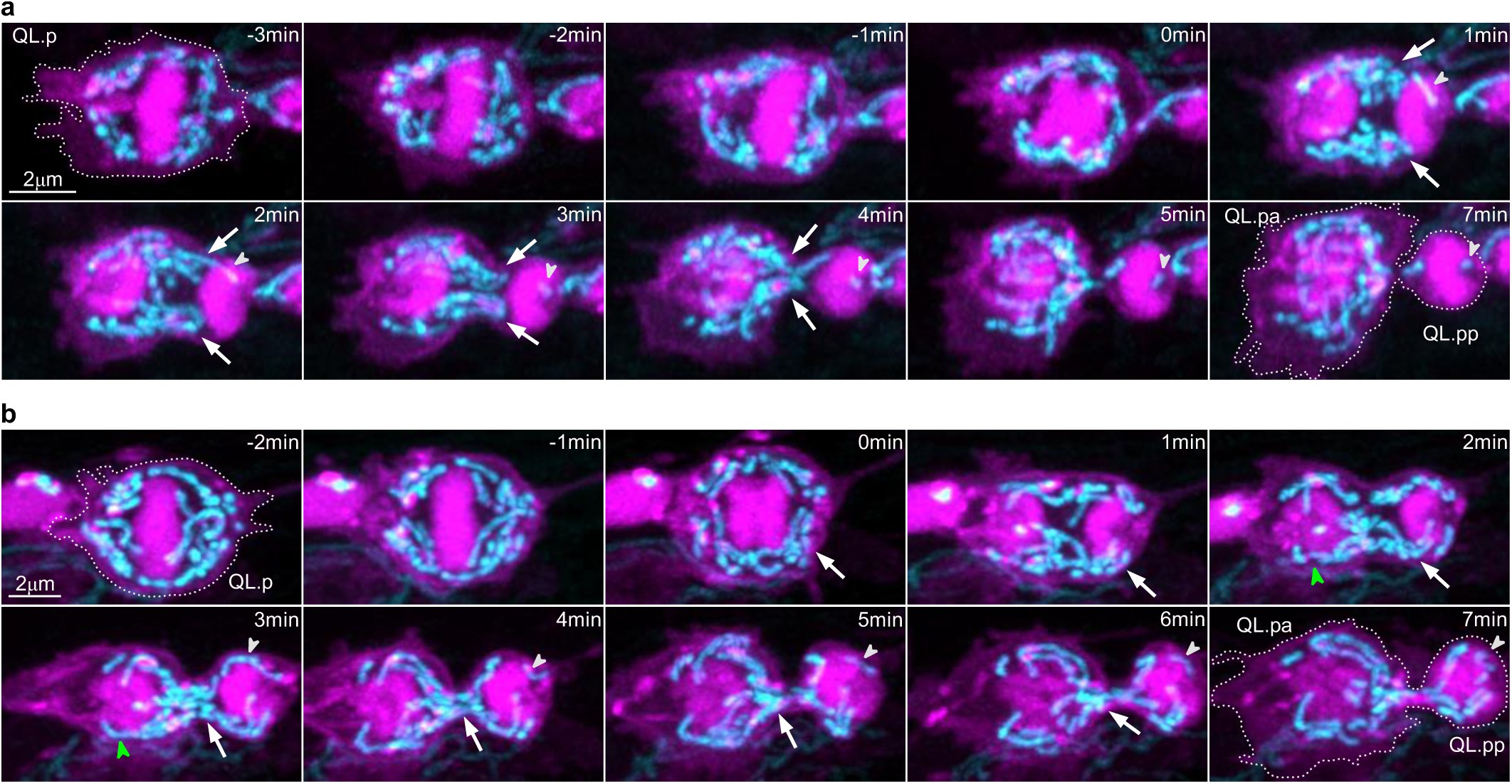
Super-resolution timeseries of mitochondrial segregation during QL.p division in wild-type animals. Super-resolution live two-colour time series of QL.p division. Plasma membrane (myristoylated mCherry) and chromatin (mCherry::his-24) are showed in magenta, mitochondria (mtGFP) in cyan (*bcIs153* transgene). Images are maximum intensity projections of aligned z-stacks. In all images, anterior is left and posterior is right. From top to bottom: wildtype examples representative of higher (**a**), and lower (**b**) mitochondrial density ratio (QL.pa/QL.pp). Arrows point to anteriorly directed transport of mitochondria. Green and white arrowheads point to mitochondrial fusion and fission, respectively. These timeseries are referred to in Figure 2.

**Figure S5.**
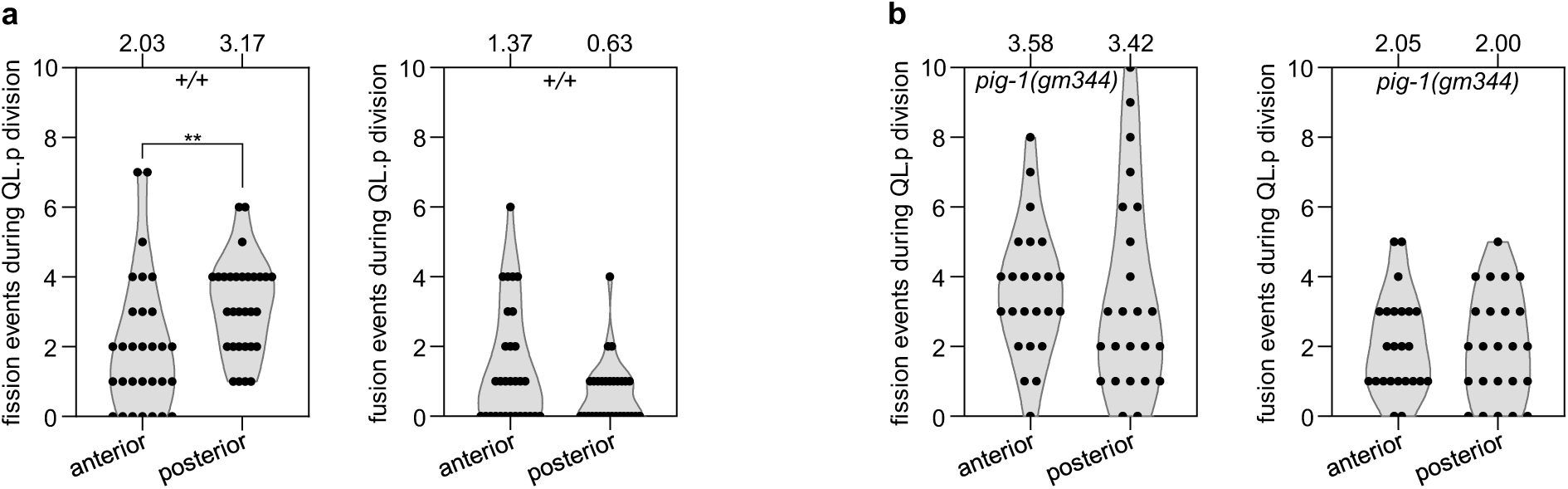
Mitochondrial fission and fusion events during QL.p division. Entire recordings were visually inspected for fission and fusion events throughout the sections of individual z-stacks. **a**,**b**) mitochondrial fission and fusion events per division in the anterior and posterior sides of QL.p between metaphase and post-cytokinesis in wildtype (*+/+*) and *pig-1(gm344)*. Statistical significances were tested performing the Fisher’s exact test. Sample sizes are 30 and 24 for *+/+* and *pig-1(gm344)*, respectively. *: P value ≤ 0.05; **: P value ≤ 0.01; ***: P value ≤ 0.001; ****: P value ≤ 0.0001. For both genotypes, animals express the *bcIs153* allele.

**Figure S6.**
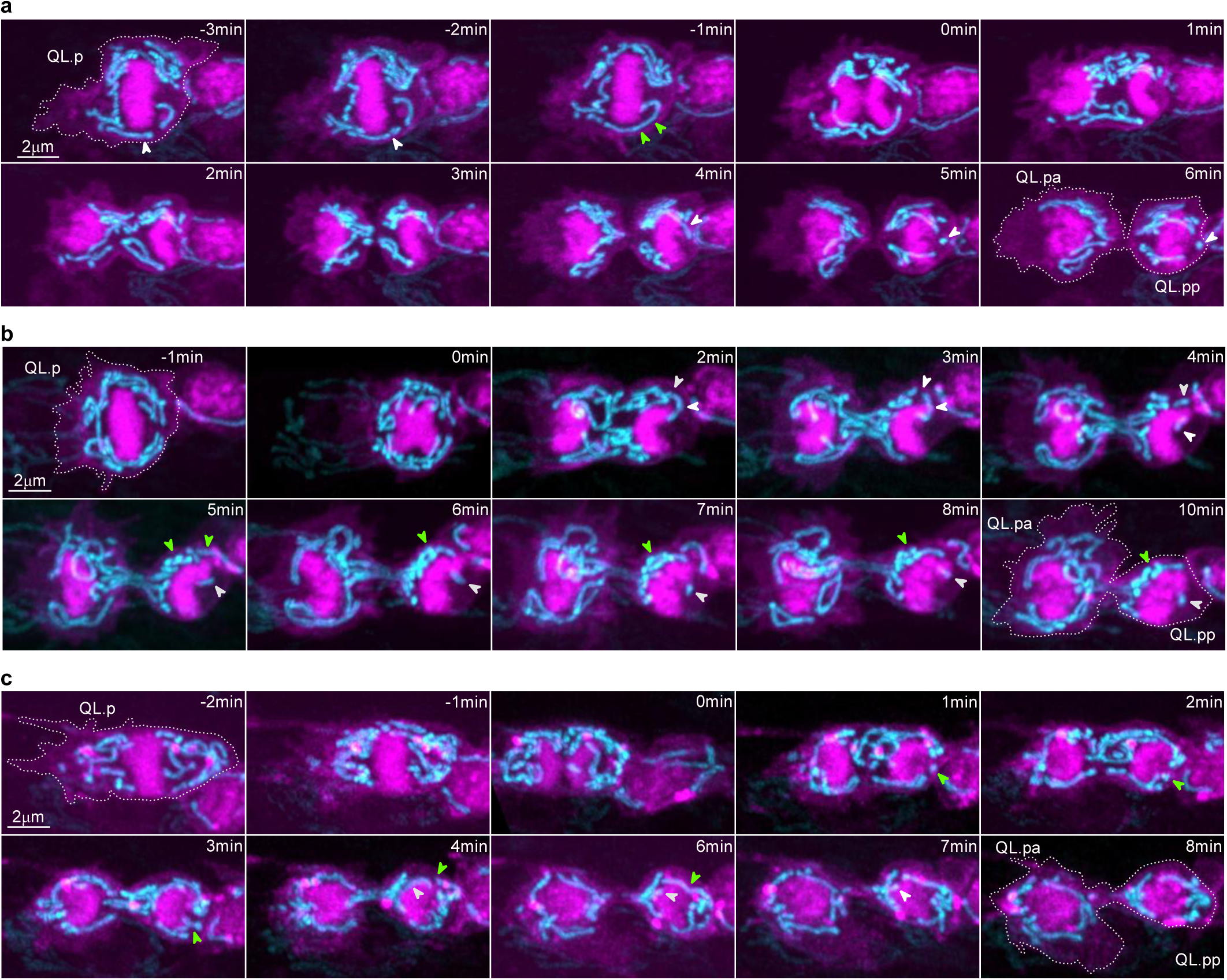
Super-resolution timeseries of mitochondrial segregation during QL.p division in *pig-1(lf)* animals. Super-resolution live two-colour time series of QL.p division. Plasma membrane (myristoylated mCherry) and chromatin (mCherry::his-24) are showed in magenta, mitochondria (mtGFP) in cyan (*bcIs153* transgene). Images are maximum intensity projections of aligned z-stacks. In all images, anterior is left and posterior is right. From top to bottom: *pig-1(gm344)* examples representative of higher (**a**), average (**b**), and lower (**c**) mitochondrial density ratio (QL.pa/QL.pp). Green and white arrowheads point to mitochondrial fusion and fission, respectively. These timeseries are referred to in Figure 2.

**Figure S7.**
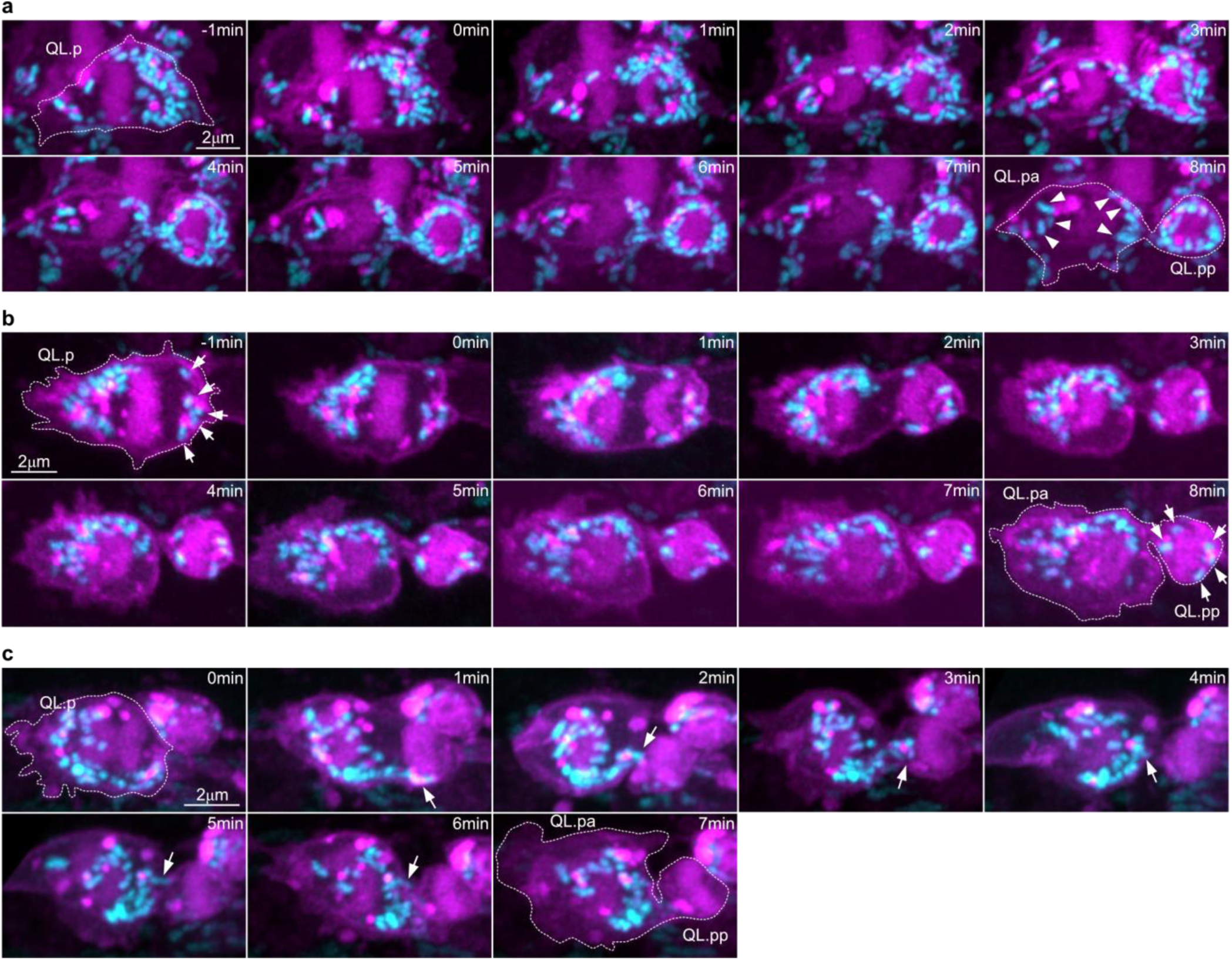
Super-resolution timeseries showing the abnormal segregation of mitochondria during QL.p division in *fzo-1(lf)* animals. Super-resolution live two-colour time series of QL.p division. Plasma membrane (myristoylated mCherry) and chromatin (mCherry::his-24) are showed in magenta, mitochondria (mtGFP) in cyan (*bcIs153* transgene). Images are maximum intensity projections of aligned z-stacks. In all images, anterior is left and posterior is right. From top to bottom: *fzo-1(tm1133)* examples representative of lower (**a**), average (**b**), and higher (+∞) (**c**) mitochondrial density ratio (QL.pa/QL.pp). Arrowheads point to QL.pa mitochondria (**a**). Arrows point to posterior mitochondria (metaphase) that were all inherited by QL.pp (**b**) and to the anteriorly directed movement of mitochondria (**c**). These timeseries are referred to in Figure 2. The example in panel S7c is not showed in Figure 2.

**Figure S8.**
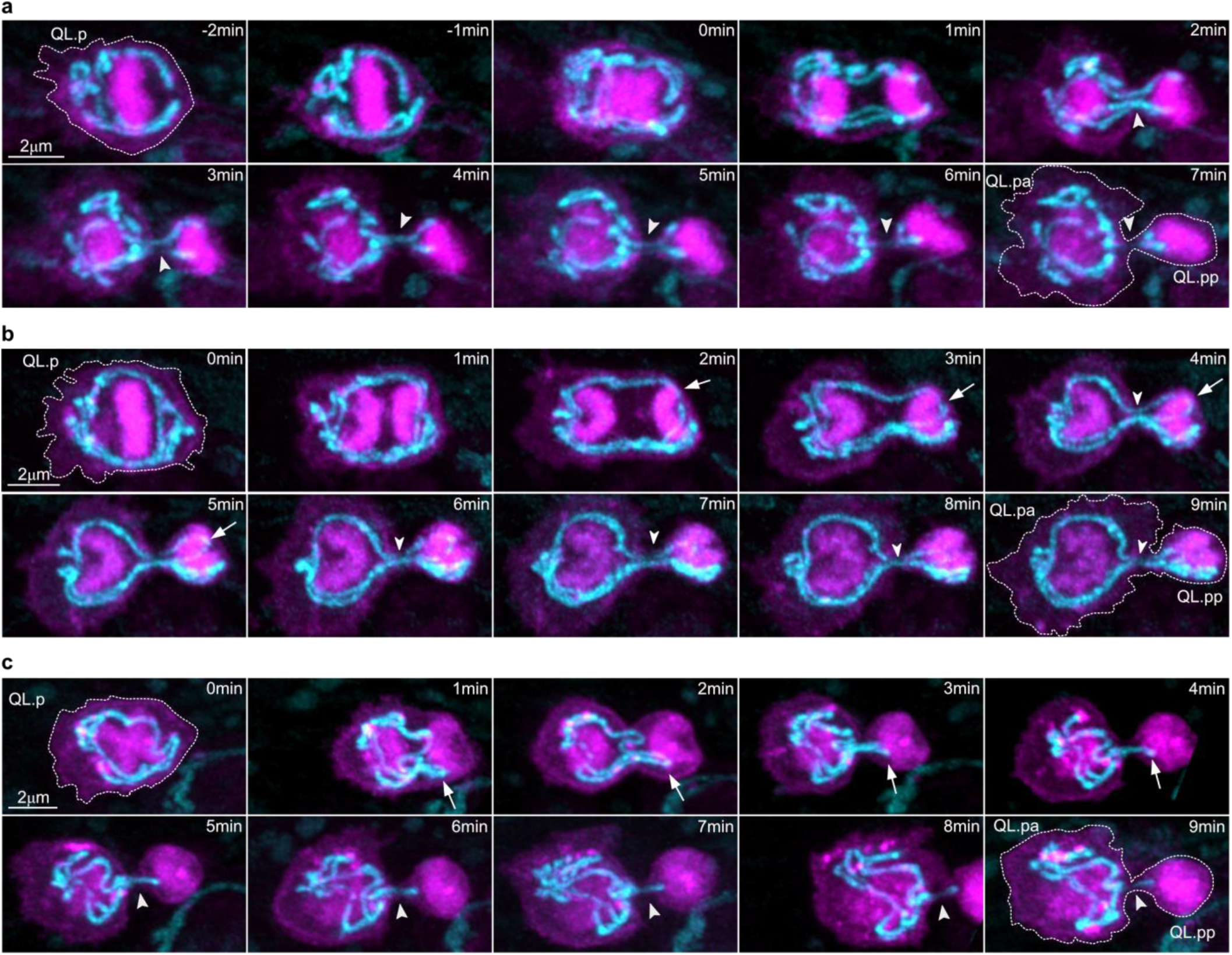
Super Resolution timeseries showing the abnormal segregation of mitochondria during QL.p division in *drp-1(lf)* animals. Super-resolution live two-colour time series of QL.p division in animals expressing *bcIs153*. Plasma membrane (myristoylated mCherry) and chromatin (mCherry::his-24) are showed in magenta, mitochondria (mtGFP) in cyan. Images are maximum intensity projections of aligned z-stacks. In all images, anterior is left and posterior is right. Top (**a**) and centre (**b**): *drp-1(bc455)* examples representative of average and lower mitochondrial density ratio (QL.pa/QL.pp). Bottom (**c**): *drp-1(tm1108)* example representative of higher mitochondrial density ratio (QL.pa/QL.pp) (see main text for further information). Arrowheads point to mitochondrial fission, whereas arrows point to a posterior mitochondrial portion looped around the posterior chromosome set blocking the anterior movement of mitochondria (**b**) and to the anteriorly directed movement of mitochondria (**c**). These timeseries are referred to in Figure 2.

**Figure S9.**
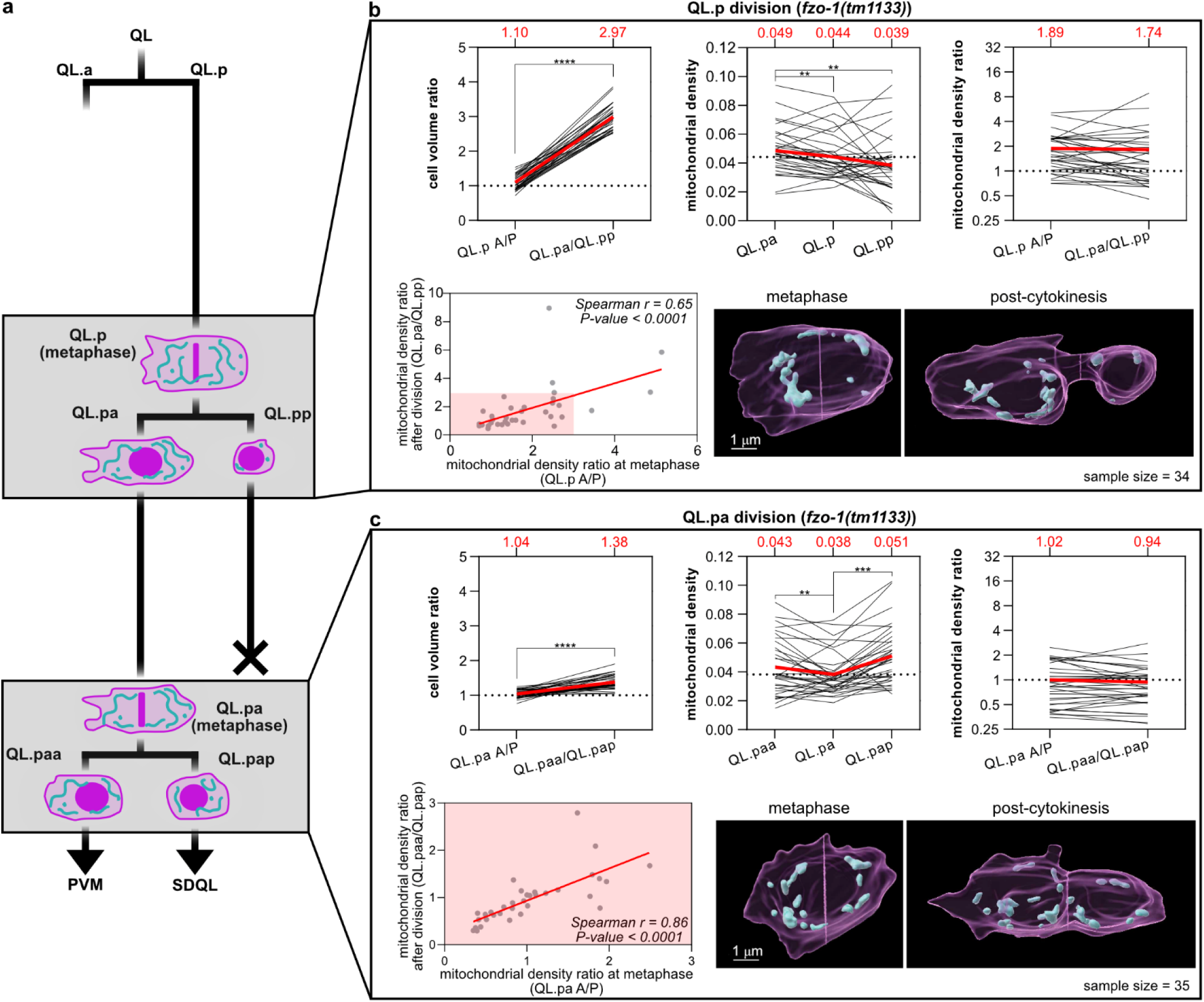
Mitochondrial partitioning in *fzo-1(lf)* mutants. **a**) Schematics of the QL.p lineage recorded in *fzo-1(tm1133)* animals expressing *bcIs153* transgenic allele. The grey boxes highlight both QL.p and QL.pa divisions. **b**) Measurements of cell volume ratio, mitochondrial density and mitochondrial density ratio during QL.p division. Top left: cell volume ratio before (QL.p A/P sides) and after division (QL.pa/QL.pp); top centre: mitochondrial density before (QL.p) and after division (QL.pa and QL.pp); top right: mitochondrial density ratio before (QL.p A/P sides) and after division (QL.pa/QL.pp); bottom left: correlation between mitochondrial density ratio before (x axes) and after (y axes) division. **c**) Measurements of cell volume ratio, mitochondrial density and mitochondrial density ratio during QL.pa division. Top left: cell volume ratio before (QL.pa A/P sides) and after division (QL.paa/QL.pap); top centre: mitochondrial density before (QL.pa) and after division (QL.paa and QL.pap); top right: mitochondrial density ratio before (QL.pa A/P sides) and after division (QL.paa/QL.pap); bottom left: correlation between mitochondrial density ratio before (x axes) and after (y axes) division; bottom right: representative 3D volumes of QL.pa division. P values are calculated using the Wilcoxon matched pairs signed rank test (both **b** and **c**, top right), the paired t-test (both **b** and **c**, top left), the RM one-way ANOVA with the Benjamini, Krieger and Yekutieli correction (**b**, top centre), the Friedman test with the Benjamini, Krieger and Yekutieli correction (**c**, top centre), and the Spearman correlation (both **b** and **c**, bottom left). Normality was tested with the Shapiro-Wilk test. *: P value ≤ 0.05; **: P value ≤ 0.01; ***: P value ≤ 0.001; ****: P value ≤ 0.0001. Red lines = mean (both **b** and **c**, top row). In the top row plots of panels **a** and **b**, individual black lines represent the trends of each division between metaphase and post-cytokinesis or between QL.p and QL.pa and QL.pp. Red lines and red numbers = average (both b and c, top row). n=35 in a and b.

**Figure S10.**
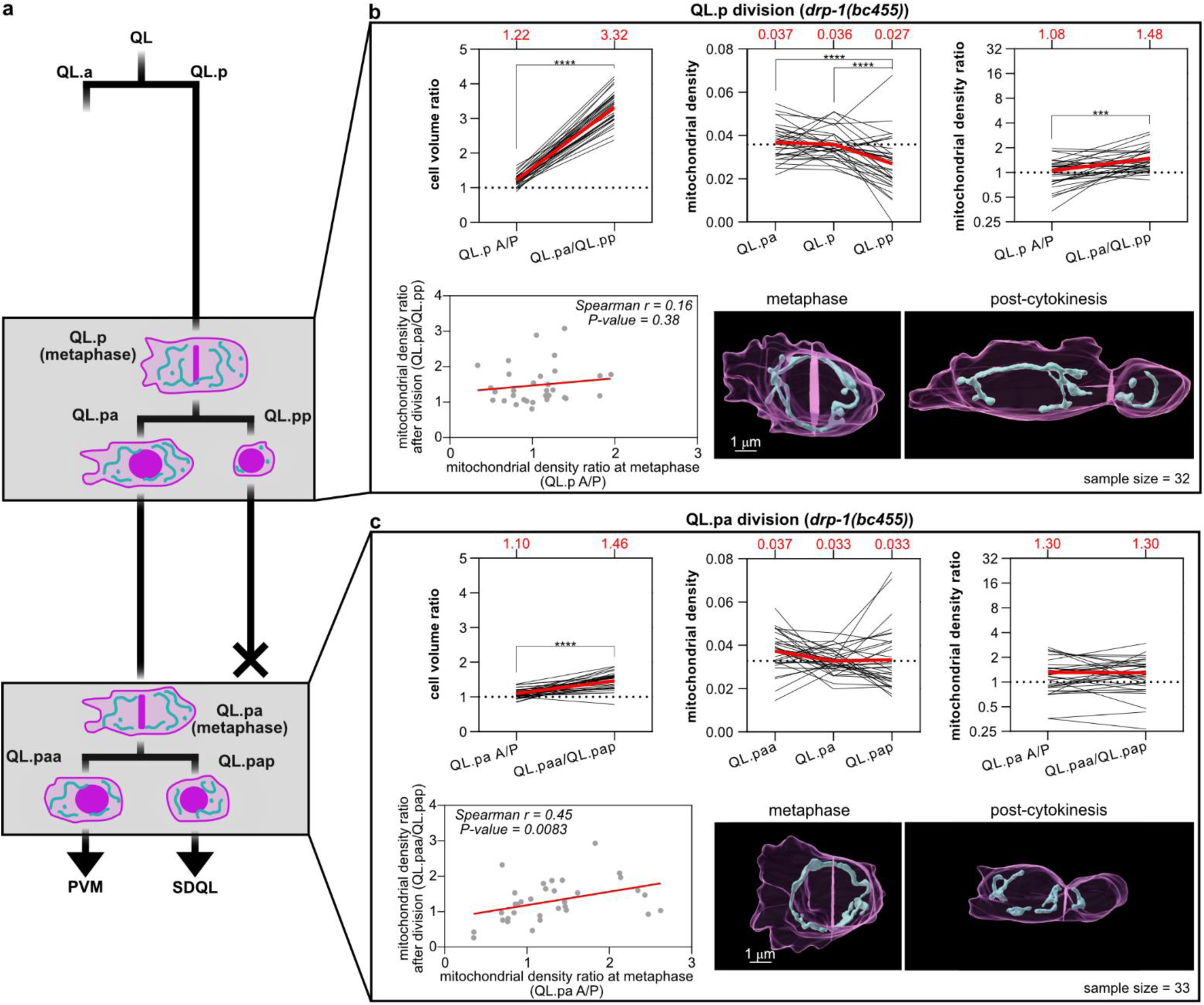
Mitochondrial partitioning in *drp-1(lf)* mutants. **a**) Schematics of the QL.p lineage recorded in *drp-1(bc455)* animals expressing *bcIs153* transgenic allele. The grey boxes highlight both QL.p and QL.pa divisions. **b,c**) Equivalent to figure S9a,b in both the arrangement and the content of plots. P values are calculated using the Wilcoxon matched pairs signed rank test (both **b** and **c**, top right), the paired t-test (both **b** and **c**, top left), the RM one-way ANOVA with the Benjamini, Krieger and Yekutieli correction (**b**, top centre), the Friedman test with the Benjamini, Krieger and Yekutieli correction (**c**, top centre), and the Spearman correlation (both **b** and **c**, bottom left). Normality was tested with the Shapiro-Wilk test. *: P value ≤ 0.05; **: P value ≤ 0.01; ***: P value ≤ 0.001; ****: P value ≤ 0.0001. Red lines = mean (both **b** and **c**, top row). In the top row plots of panels **a** and **b**, individual black lines represent the trends of each division between metaphase and post-cytokinesis or between QL.p and QL.pa and QL.pp. Red lines and red numbers = average (both b and c, top row). n=35 in a and b.

**Figure S11.**
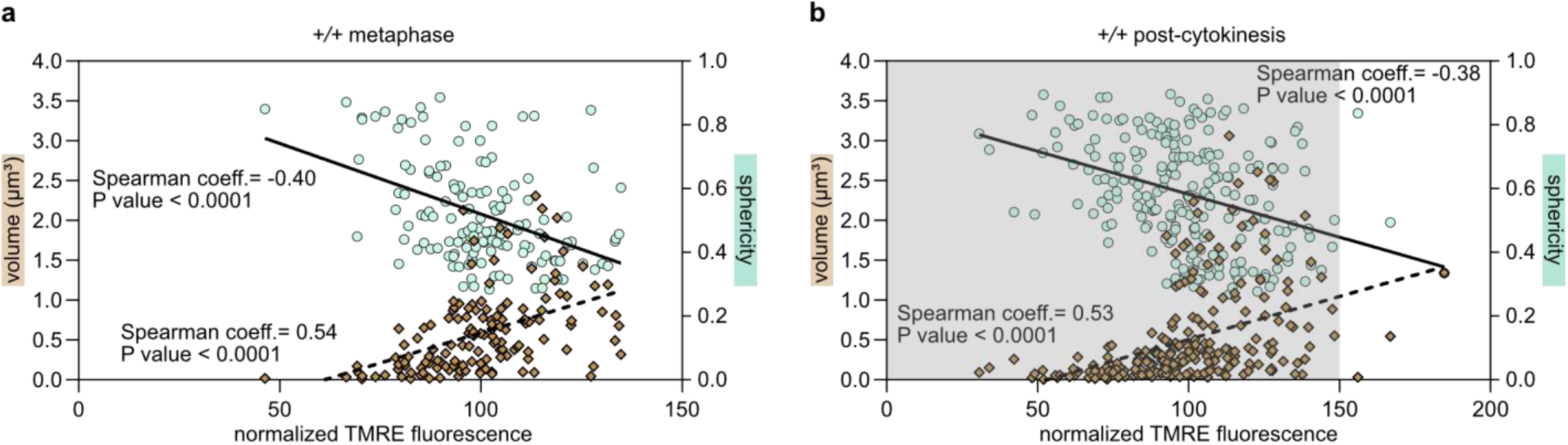
Mitochondria activity (TMRE) and mitochondrial morphology are correlated. Correlation between normalized TMRE fluorescence and mitochondrial sphericity and volume at single mitochondrion level at metaphase (**a**) and post-cytokinesis (**b**), in wildtype animals expressing the *bcIs158* allele. The mean TMRE intensity within each mitochondrion was normalized by dividing it with the mean TMRE intensity of all mitochondria in the respective cell (QL.p for metaphase or QL.pa and QL.pp, together, for post-cytokinesis) (see methods for further information). The grey panel in **b** refers to the XY dimensions of in **a**. Continuous line= linear regression of the TMRE-mitochondrial sphericity correlation; dotted lines= linear regression of TMRE-mitochondrial volume correlation.

**Figure S12.**
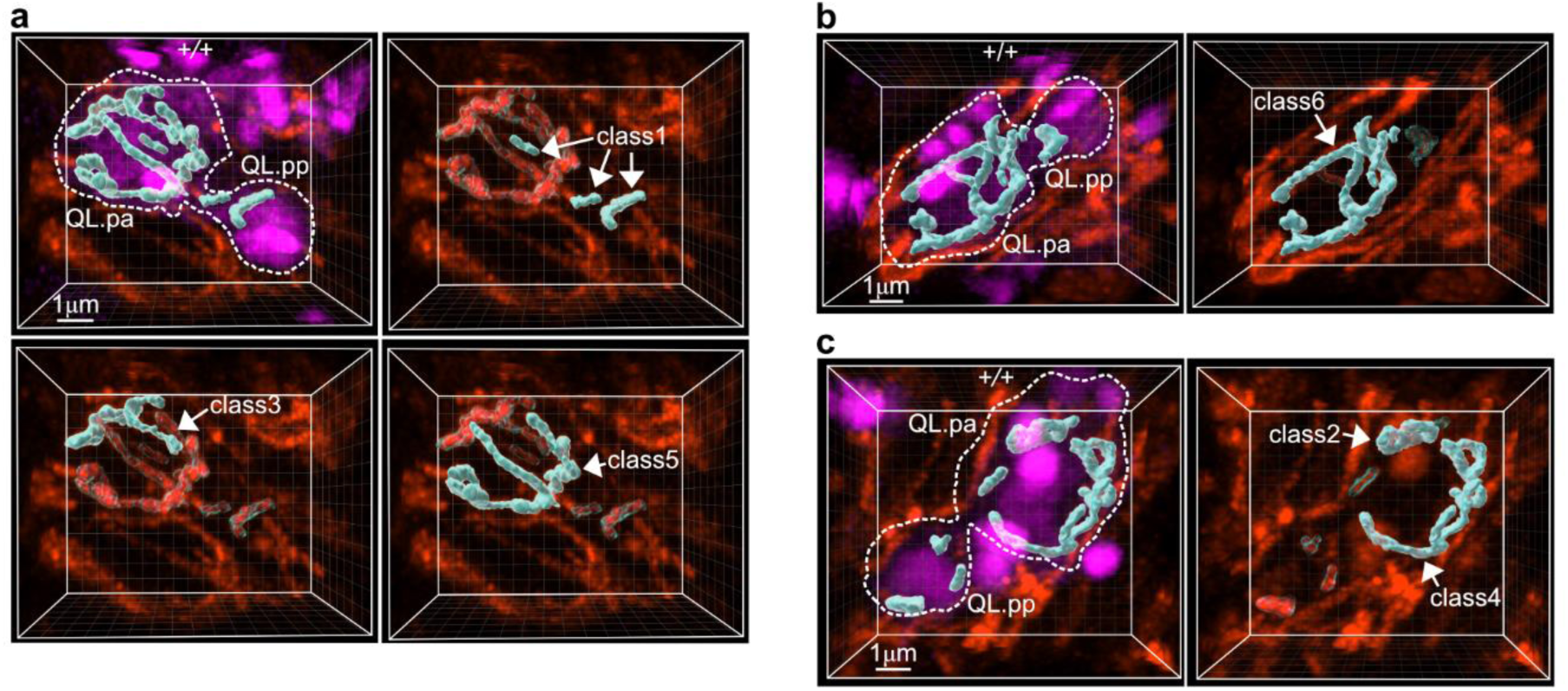
Mitochondrial classes determined to measure TMRE in individual organelles. Representative images showing examples of the six volume classes of mitochondria used to study TMRE fluorescence intensity in relation to mitochondrial morphology (mitochondria are bigger and longer and/or branched going from classes 1 to 6) in animals expressing the *bcIs158* allele. The six classes span equally the wildtype volume range 0.0043µm^3^-3.0620 µm^3^. **a**) mitochondrial classes 1,3 and 5; **b**) mitochondrial class 6; **c**) mitochondrial classes 2 and 4.

**Figure S13.**
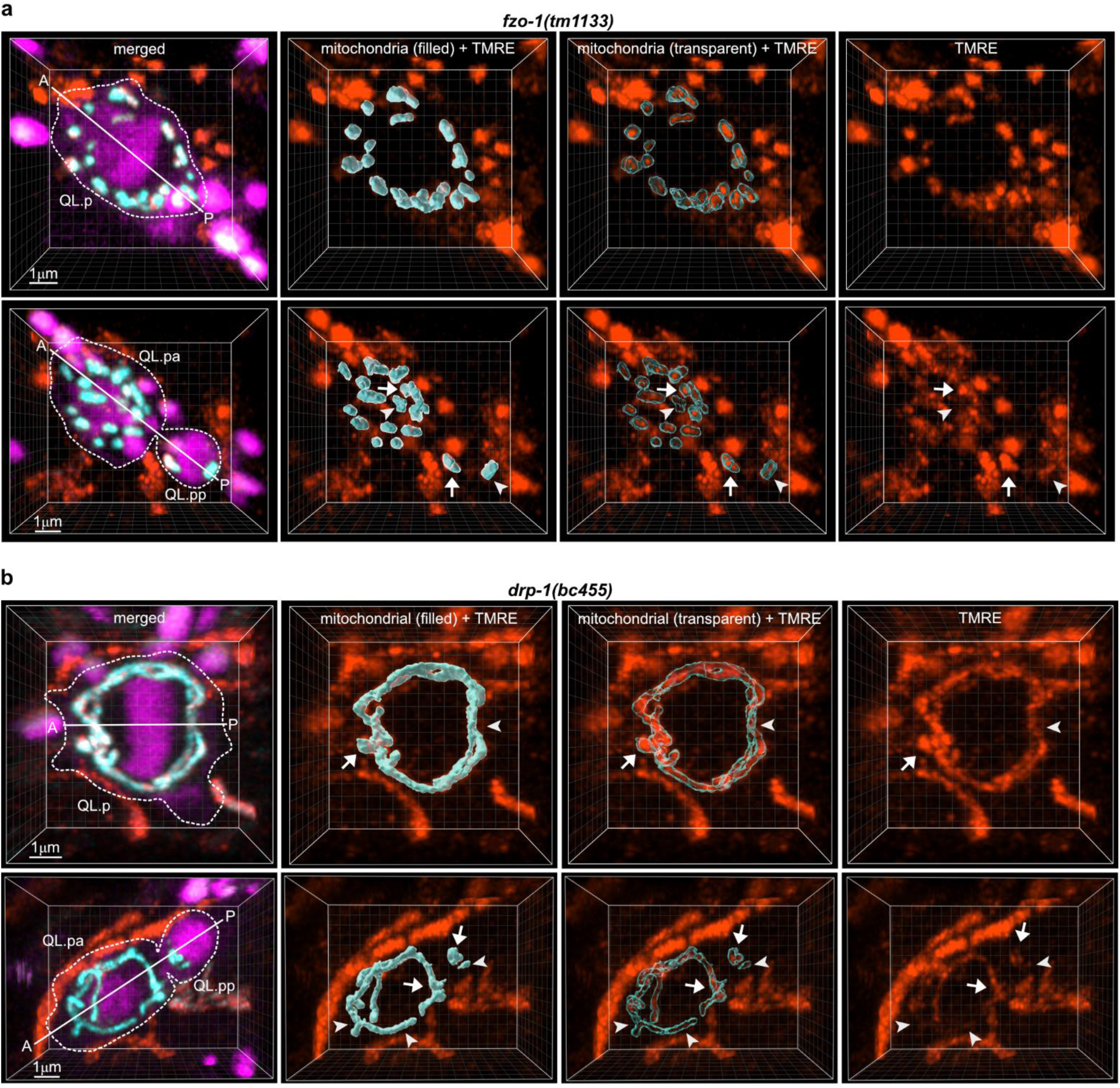
Mitochondria activity (TMRE) during QL.p division in *fzo-1(lf)* and *drp-1(lf)* mutants. **a**) representative 3D image of *fzo-1(tm1133)* (**a**) and *drp-1(bc455)* (**b**) QL.p cells labelled with TMRE at metaphase (top tow) and post-cytokinesis (bottom row). In both top and bottom left images (merged): both myristoylated SFmTurquoise2ox (cell membrane) and SFmTurquoise2ox::his-24 (chromatin) are shown in magenta, mtGFP (mitochondria) in cyan, and TMRE in orange. The A-P axes show the orientation of QL.p division. Arrows and arrowheads point respectively to mitochondria with higher and lower TMRE intensities. TMRE fluorescence intensities are measured within each volume of mitochondria (see “mitochondria (3D) + TMRE” images) in Imaris. For both genotypes, animals express the *bcIs158* allele.

**Figure S14.**
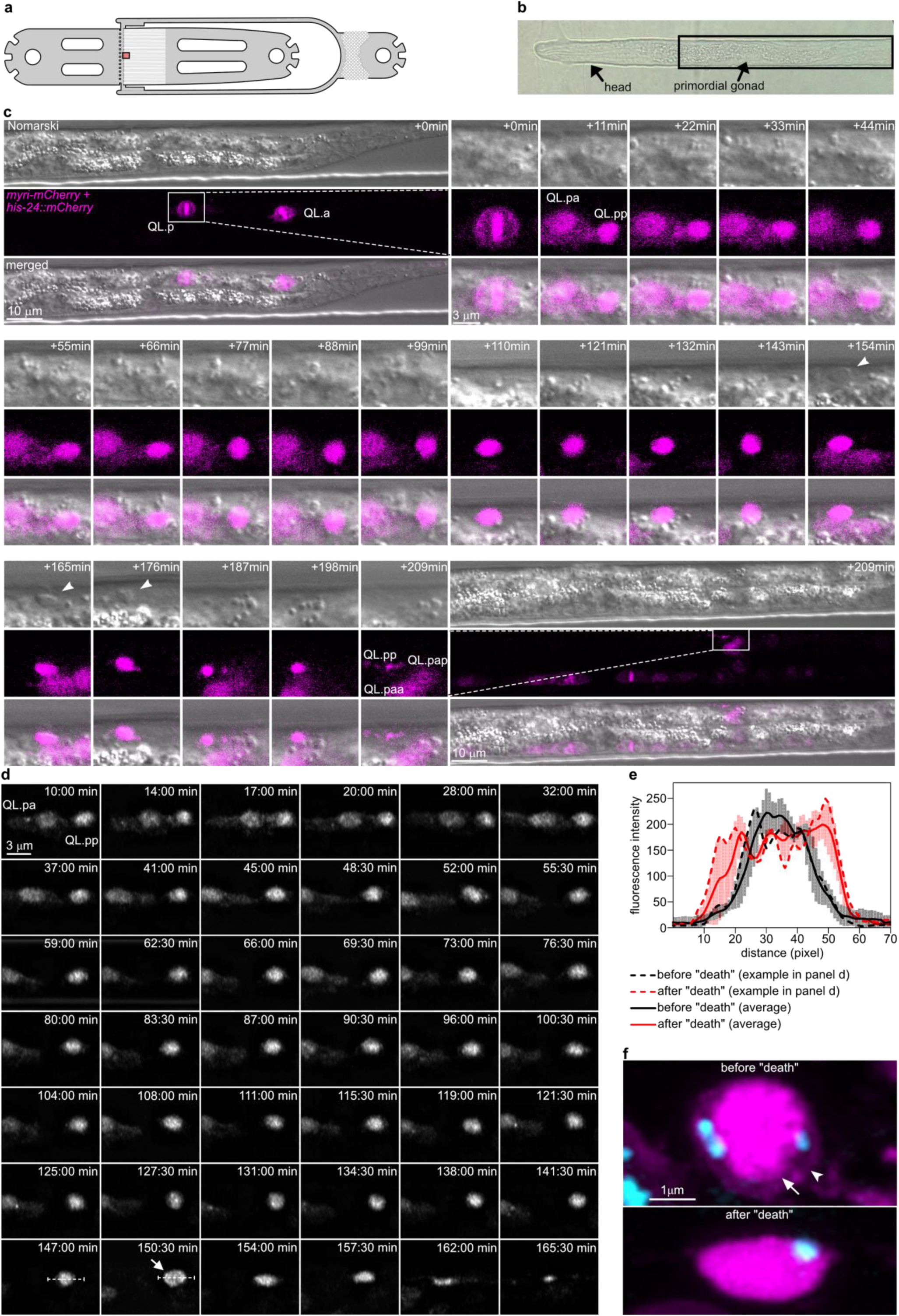
(previous page). Determination of QL.pp survival time in fluorescence microscopy images. **a**) schematics of the microfluidics device used in this study. The red box refers to **b**. **b**) L1 larva correctly trapped in a microfluidics channel. The rectangular highlight represents the field of view used for both Nomarski + LSM confocal imaging (**c**) and Airyscan multiplex confocal imaging (**d**). **c**) Nomarsky and LSM microscopy of QL.p lineage. The top left and bottom right images represent the enlarged fields of view referring to the first and last time point of the timeseries, respectively. For all images, top, center and bottom represent Nomarsky microscopy, confocal microscopy, and merged images. Anterior and posterior are on the left- and right-hand sides, respectively. Between time points +11 and +264, the time-series shows alternate images of the original time series. White arrowheads indicate the refractile appearance of the cell corpse. **d**) Airyscan multiplex confocal recording of the QL.pp cell death process. The white arrow indicates the first appearance of QL.pp as an elliptical object, while the dotted line is the segment of the image used to construct the fluorescence profiles given in **e**. **e**) Fluorescence intensity profiles of QL.pp mCherry chromatin and cell membrane markers. The continuous trends represent the average of 4 cells, while error bars represent the standard deviation. **f**) Airyscan Super Resolution images of QL.pp before (top) and after (bottom) the morphological change used as the cell death time point in this study (anterior and posterior are left and right, respectively). Before “death”, QL.pp is round and its cell membrane (arrowhead) and nucleus (arrow) are distinguishable, whereas after “death” the nucleus is disintegrated and the release of mCherry::his-24 homogenizes the fluorescence intensity throughout the whole corpse. During “death”, the QL.pp round shape turns into a distinct ellipsoid having the longer axes always oriented along the anterior-posterior axes of the animal. In this Figure, animals express the *bcIs153* allele.

**Figure S15.**
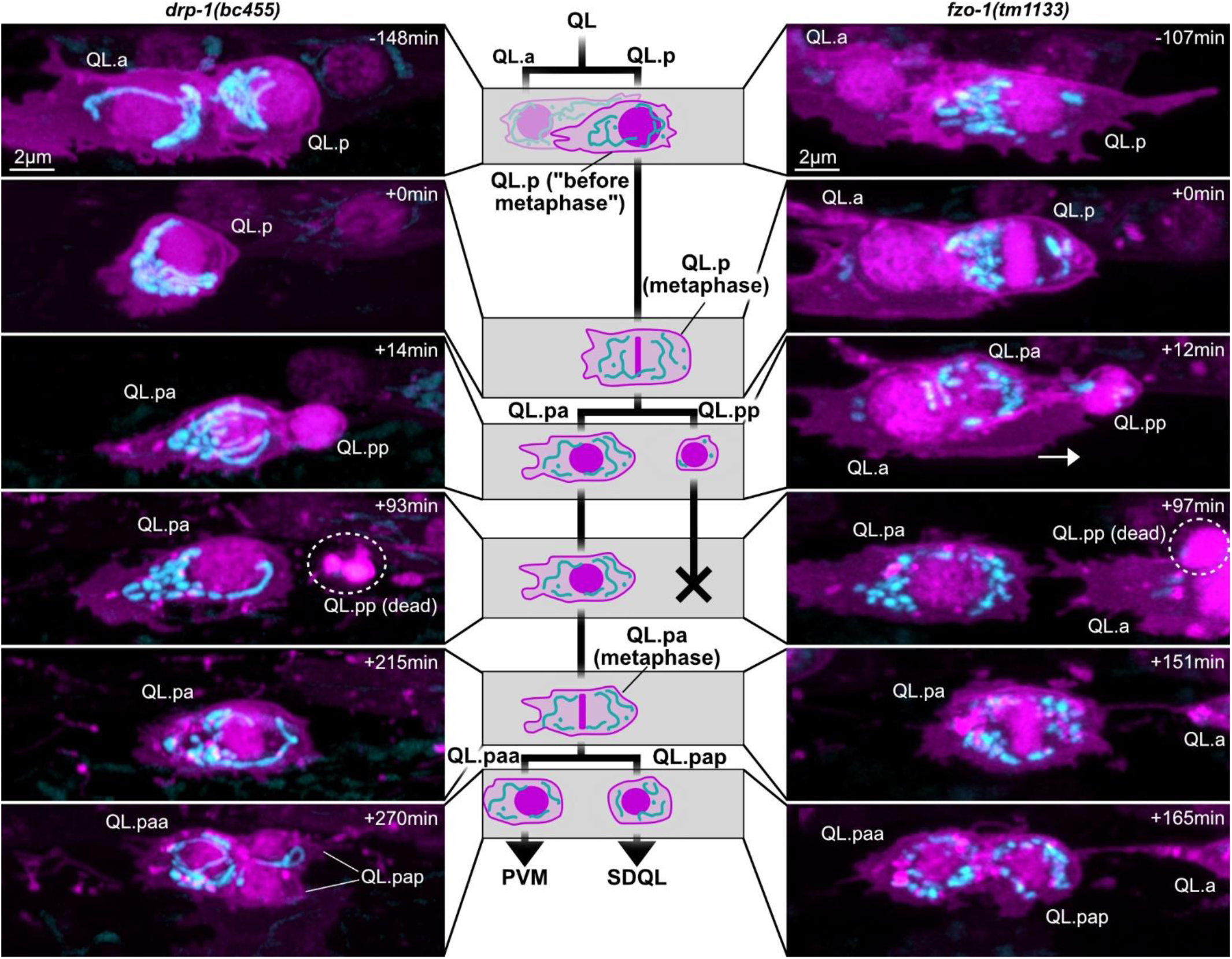
In absence of mitochondrial dynamics, QL.pp can inherit few or no mitochondria. Super-resolution live two-colour images of QL.p lineages. Plasma membrane (myristoylated mCherry) and chromatin (mCherry::his-24) are showed in magenta, mitochondria (mtGFP) in cyan (*bcIs153* allele). Images are maximum intensity projections of aligned z-stacks. Both image columns represent examples of abnormal QL.p mitochondrial partitioning in *drp-1(bc455)* and *fzo-1(tm1133)* (left and right, respectively) that caused QL.pp to inherit no mitochondria (left, time +14) or few mitochondria (right, time +12). In both cases, QL.pp died relatively fast (dotted circles) (+93 and +97 for *drp-1(bc455)* and *fzo-1(tm1133)*, respectively) (compare with QL.pp death time ranges in Fig.4 **e** and **g**). All images refer to the relative time point depicted along the schematics of the QL.p lineage shown in the centre. L1 larvae developed in the microfluidics device at high resolution and images were taken in Airyscan super resolution. Images were taken in super resolution even before QL.p division and during the disappearance of QL.pp cells in these abnormal cases for illustration (see also the abnormal distribution of mitochondria in both QL.p examples soon after QL division). The time scale is centred to metaphase (time +0min).

**Figure S16.**
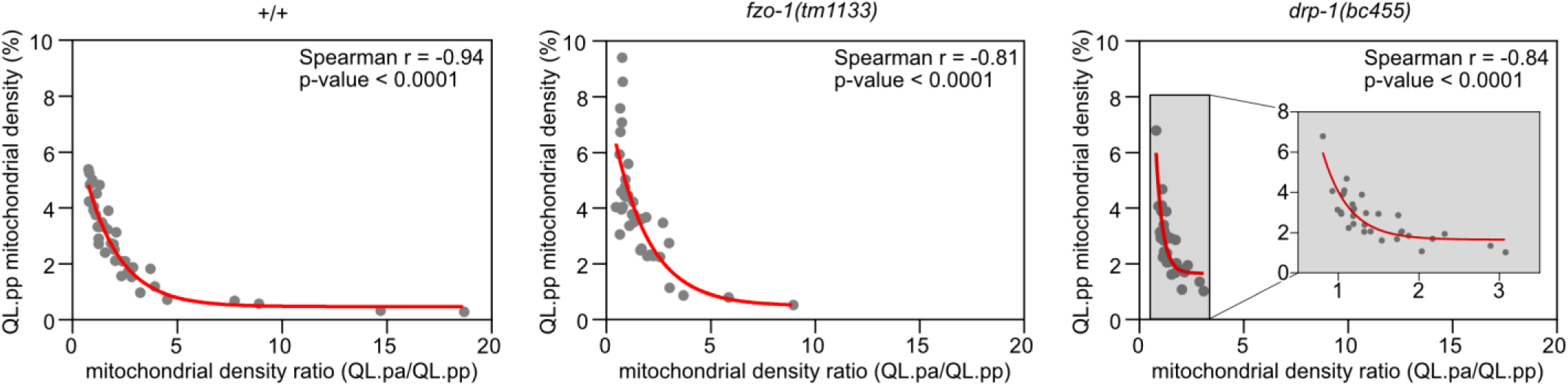
The mitochondrial density ratio and the mitochondrial density of QL.pp are nonlinearly correlated. correlation between QL.pp mitochondrial density (%) and the mitochondrial density ratio (QL.pa/QL.pp) in wildtype (*+/+*) (left), *fzo-1(tm1133)* (centre), and *drp-1(bc455)* (right). Red lines = monoexponential decay regression fitted on scattered plots. All correlations were analysed calculating the Spearman correlation coefficient *r* and its related p-value. For all genotypes, animals express the *bcIs153* allele.

